# Sensitive and Selective Polymer Condensation at Membrane Surface Driven by Positive Co-operativity

**DOI:** 10.1101/2022.07.20.500707

**Authors:** Zhuang Liu, Arun Yethiraj, Qiang Cui

## Abstract

Biomolecular phase separation has emerged as an essential mechanism for cellular organization. How cells respond to environmental stimuli in a robust and sensitive manner to build functional condensates at the proper time and location is only starting to be understood. Recently, lipid membranes have been recognized as an important regulatory center for biomolecular condensation. However, how the interplay between the phase behaviors of cellular membranes and surface biopolymers may contribute to the regulation of surface condensation remains to be elucidated. Using simulations and a mean-field theoretical model, we show that two key factors are the membrane’s tendency to phase separate and the surface polymer’s ability to reorganize local membrane composition. Surface condensate forms with high sensitivity and selectivity in response to features of biopolymer when positive co-operativity is established between coupled growth of the condensate and local lipid domains. This effect relating the degree of membrane-surface polymer co-operativity and condensate property regulation is shown to be robust by different ways of tuning the co-operativity, such as varying membrane protein obstacle concentration, lipid composition and the affinity between lipid and polymer. The general physical principle emerged from the current analysis may have implications in other biological processes and beyond.

---

Cells are compartmentalized into distinct functional regions often surrounded by biological membranes, known as organelles, for carrying out the diverse biochemistry of life. In addition, phase separation driven by weak, multivalent interactions among biomolecules has emerged as an essential mechanism for cellular compartmentalization (1). The biomolecular condensates formed through phase separation, enriched in selected proteins and often RNAs, are known as membraneless organelles (MLOs) and have been revealed to play an essential role in cell physiology (2–6). Unlike membrane-bound organelles, the formation of these phase-separated condensates are typically reversible in response to cellular cues (7–12). The mechanism of how cells respond to stimuli in a robust and sensitive manner to build specific functional condensates in a spatially and temporally relevant manner is only starting to be understood.

In recent years, biological membranes have been recognized as a key regulatory center for controlled condensate formation in cells (13–17). In particular, prewetting appears to be a mechanism by which cells could exert spatiotemporal control over the assembly of biomolecular condensates (18– 20). In general, proteins separate into a coexisting dense phase (condensate) and a dilute phase in the cytoplasm when their mutual interaction strength reaches a certain threshold value *J*_*c*_. In a prewetting process, proteins (or biopolymers in general) are attracted to a surface, which enables their condensation at interaction strengths lower than *J*_*c*_. This surface condensate (prewetting phase) is restricted only to the vicinity of the surface, as bulk condensation is not favorable. Yet beyond merely serving as an attractive two-dimensional surface, biological membranes are fluidic structures with heterogeneous and complex lipid and protein compositions that can phase separate on their own (21–24). The goal of this work is to understand the interplay between the phase behaviors of biological membranes and the biopolymers at the membrane surfaces. We aim to establish general principles that might govern how cells regulate surface condensation.

In an interesting study(18), Machta and co-workers investigated the coupled phase behavior of a 2-component membrane and polymers; they focused on the conditions of coexistence of different surface phases and presented a framework for analyzing the problem through simulation and theory. The main conclusion of their work was that proximity to the membrane critical point greatly enhances condensate formation. An important feature of biological membranes that their work did not include is the presence of membrane proteins. Various experimental and theoretical studies have estimated that the area fraction of proteins in biological membranes ranges from 20 to 75% depending on the membrane type (25–31), and that these embedded “protein obstacles” have significant influences on the phase behaviors of the surrounding lipids (32–34). Therefore, a realistic model for studying the coupled phase behaviors of membrane and surface biopolymers must take the effect of protein obstacles into account (Fig. 1). Furthermore, as the functions of biomolecular condensates are dependent on their biophysical properties (35–43), it’s critical to understand how the phase behaviors of membranes and surface biopolymers regulate the condensate properties beyond the condition of formation; this topic has been relatively unexplored in previous studies (18, 19, 44, 45).

**Fig. 1.**
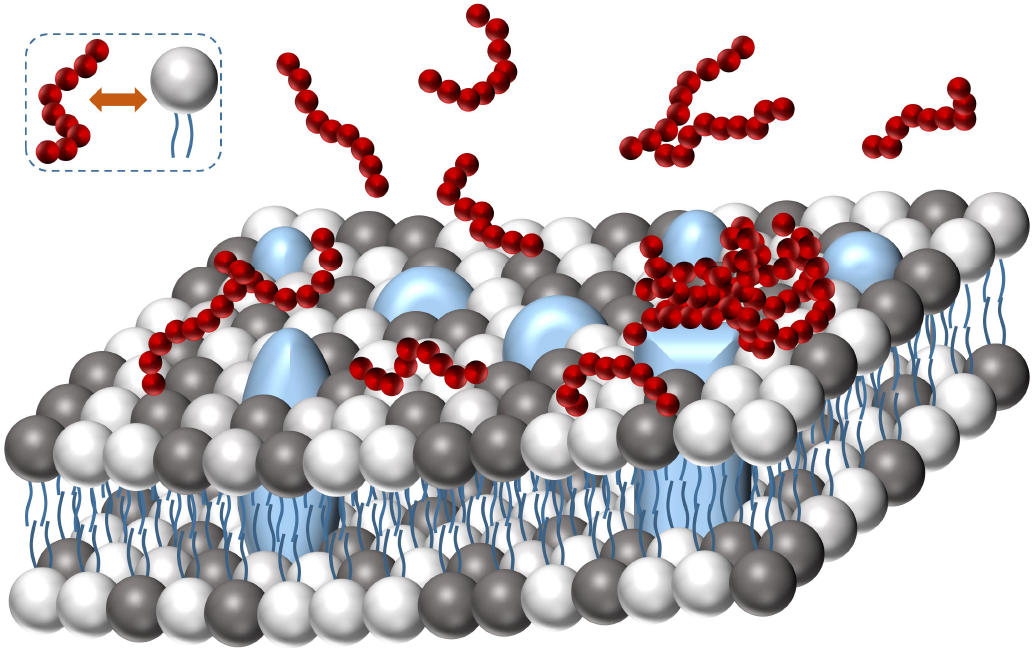
Schematic of biopolymers (red) at the surface of a membrane consisting of two types of lipids (black and white) and embedded protein obstacles (blue), where biopolymers are attracted to one type of lipid (white). We investigate here how obstacles and the variation of other parameters of the system (e.g., lipid composition *f*_*A*_; lipid-lipid interaction strength *J*_*m*_; lipid-polymer interaction strength *h*_*t*_ and range *l*) modify the coupled phase behavior of biopolymers and lipids to regulate surface condensation.

In this work, we first explore the effect of membrane protein obstacles on regulating surface condensation through grand canonical Monte Carlo (GCMC) simulations and a mean-field theory (MFT). Our simulation results show that the presence of protein obstacles in the membrane at physiological con-centrations enhances the sensitivity and selectivity of surface condensation and membrane reorganization to the property of the biopolymer, and that such effect is observed over a range of membrane conditions. Furthermore, our theoretical analysis confirms the findings of the simulations, and reveals that such obstacle effect originates from the positive co-operativity between the coupled growth of local lipid domains and surface condensates. The general significance of membrane-surface polymer co-operativity to surface condensation is then further verified through simulation and theoretical analyses of several other model systems in which the degree of membrane-surface polymer co-operativity is perturbed in distinct ways. Although the simple models used here do not represent the rich membrane chemistry present in biology, the underlying physical principles are robust and potentially relevant to many biological processes.

## Simulation Results

### Simulation Model for the Effect of Protein Obstacles on Membrane Phase Behaviors

Before exploring the effect of obstacles on surface condensation, we first briefly review the basic simulation framework and the effect of obstacles on the phase behavior of membrane. We model binary lipid membrane embedded with protein obstacles using a fixed composition 2D Ising model on square lattice with an attractive/repulsive nearest neighbor (NN) interaction energy *J*_*m*_(*k*_*B*_*T*) between like/unlike lipid pairs (see *SIAppendix*, Fig. S1 A). The Hamiltonian of the system is

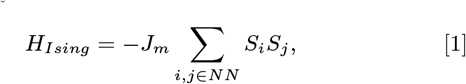

where *i* and *j* label lattice sites, *S*_*i*_ = 1*/* − 1 for lipid species A/B and *S*_*i*_ = 0 if it’s occupied by an obstacle, indicating that obstacles are inert and have no (preferential) interaction with either lipid. In the simulations, *J*_*m*_ and the fraction of obstacle sites (*f*_*o*_) are varied to describe different membrane conditions, while the number of A lipid is always kept equal to that of B (*M* =∑_*i*_ *S*_*i*_ = 0), representing fixed lipid compositions. Furthermore, we consider two types of obstacles: immobile and mobile (floating). Immobile obstacles describe integral membrane proteins attached to the cytoskeleton or very large integral membrane proteins that diffuse much slower than lipids, while mobile obstacles describe unattached floating membrane proteins. In a simulation with immobile obstacles, the positions of obstacle sites are fixed at initialization and the only MC move is the swap of unlike lipid pairs, while in simulations with mobile obstacles, swap moves between lipids and obstacles are included. We refer to immobile/mobile obstacles as obstacles/floating obstacles hereafter for simplicity. Our simulations show that the presence of obstacles significantly suppresses lipid phase separation, in agreement with previous studies (32–34) (see *SIAppendix* and Fig. S1-3 for detailed discussions). For example, as the fraction of obstacles increases from 0 to 0.1 and 0.3, the critical membrane coupling 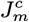 increases from 0.35 *k*_*B*_*T* to 0.4 and 0.7 *k*_*B*_*T*, respectively (Fig. S1C-E). The effect of floating obstacles, however, is far less prominent (Fig. S2).

### Membrane obstacles enhance the sensitivity and selectivity of surface condensation

We next explore the effect of membrane obstacles on surface condensation by coupling the Ising membrane with lattice polymers in our simulation (Fig. 2C). Specifically, the Ising membrane is simulated as described in the previous section and placed at the bottom of the simulation box (*z* = 0). Lattice polymers are 9 monomers long with attractive NN interaction energy *J*_*p*_(*k*_*B*_*T*), and are kept at fixed chemical potential following the conventional GCMC algorithm (46). Following Rouches et al. (18), the polymers are coupled to the membrane through tethers, which are attached only to lipid species A and extend 5 lattice sites straight into the bulk. Tethers form favorable interactions with polymers, and can translate on the 2D membrane. It should be noted that the inclusion of tethers here is not essential, and is merely one way of implementing attraction between polymers and selected lipid species, without which the membrane phase behavior does not affect surface condensation. The major conclusions from this study are not subject to specific forms of polymer-lipid coupling and apply more generally, as further discussed in the section “*General Significance of Membrane-surface polymer Co-operativity to Surface Condensation*”.

**Fig. 2.**
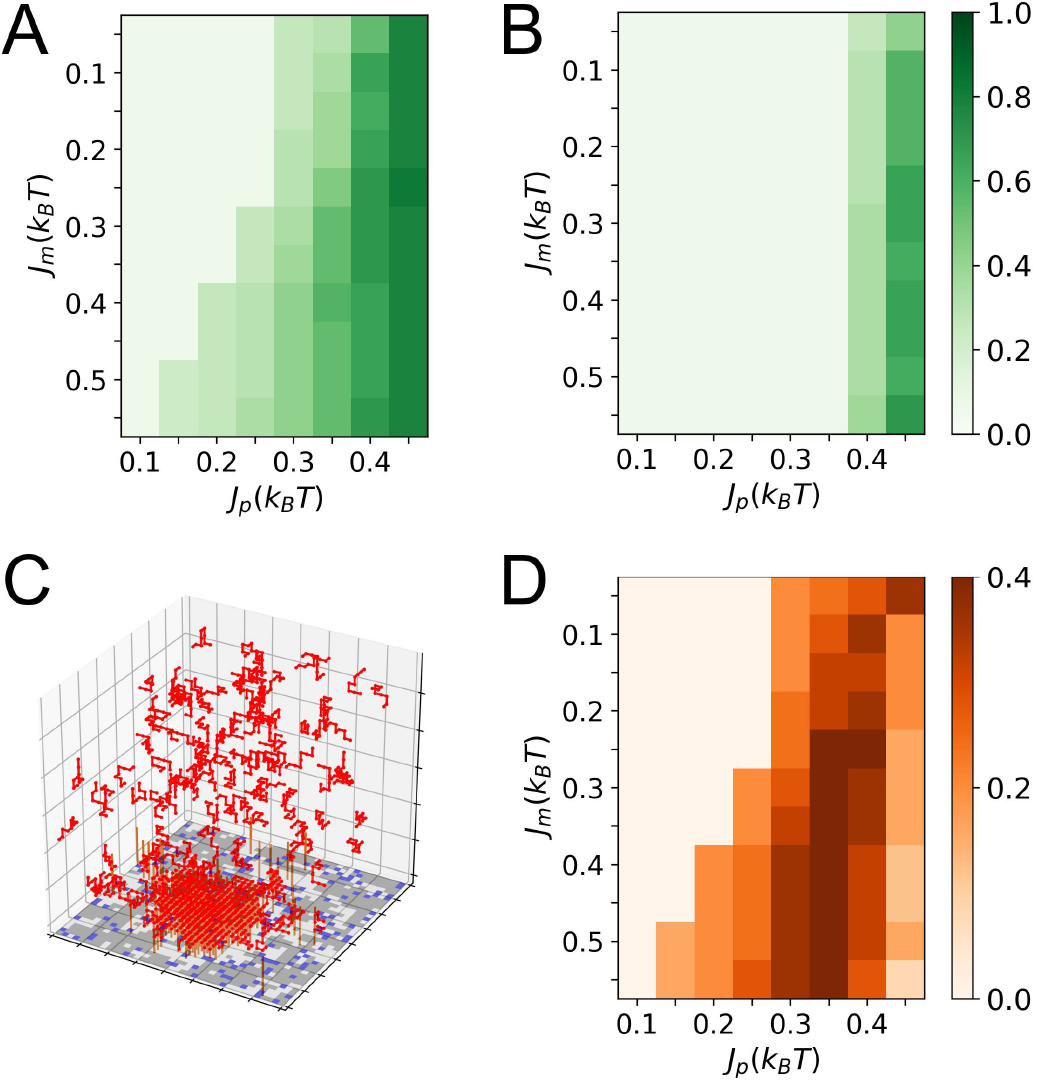
Effect of obstacles on the densities of surface condensates. Polymer density in surface condensates formed at different *J*_*p*_(*k*_*B*_*T*) and *J*_*m*_(*k*_*B*_*T*) without obstacles (A); and with obstacles at area fractions of 0.3 (B). (C) Snapshot of a simulation with *J*_*p*_ = 0.45 *k*_*B*_*T* and *J*_*m*_ = 0.35 *k*_*B*_*T* with floating obstacles at an area fraction of 0.1. Red chains represent polymers, white, grey and blue squares on the bottom plane represent the two lipid components A and B and protein obstacles of the membrane. Brown straight chains sticking out of the bottom plane represent tethers that only connect to white lipids and have a favorable interaction with the red polymers. (D) Polymer density differences between (A) and (B). Polymer density is calculated as the fraction of lattice sites occupied by red polymers in a 5 *×* 5 *×* 5 surface region. The low polymer densities (<0.1) in (A) and (B), (e.g., the columns of *J*_*p*_ = 0.1 *k*_*B*_*T*) represent the density of dilute surface phase before prewetting transitions. The tether density of the membrane is 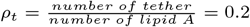 in all cases.

Without the membrane, the lattice polymers can phase separate in the bulk when *J*_*p*_ is increased to 0.55 *k*_*B*_*T* (see *SIAppendix*, Fig. S4A and B), and thus we focus on *J*_*p*_ < 0.55 *k*_*B*_*T*, where the surface condensates formed are prewetting phases. Typical surface condensates formed in prewetting are shown in the simulation snapshots in Fig. 2C and *SIAppendix* Fig. S4C, which are thin layers of polymer aggregates (see *SIAppendix* Fig. S5D). Indeed, when *J*_*p*_ reaches 0.55 *k*_*B*_*T*, surface condensates start to grow into the bulk (see *SIAppendix*, Fig. S4D).

Next, we analyze how the assembly of surface condensate responds to the change of *J*_*p*_ under different membrane conditions. The change of *J*_*p*_ is meant to be a simple model for the variation of biopolymer sequence, the state of posttranslational modification (e.g., phosphorylation) or local solution condition (e.g., pH), which are regulatory mechanisms that cells exploit to control biomolecular phase separation (47).

As shown in Fig. 2A, without obstacles, the density of the surface condensate formed increases with both *J*_*m*_ and *J*_*p*_, as manifested by the deeper color towards the lower right of the heat map, whereas, at high *J*_*p*_ values, the dependence on *J*_*m*_ is significantly weakened. Such qualitative trends remain in the corresponding heat maps after introducing obstacles (*Fig. 2B* and *SIAppendix* Fig. S5B and C), while high condensate densities are observed at higher *J*_*p*_ values as the amount of obstacles increases. It should be emphasized that high density of phase separating biomolecules in condensate relative to the dilute surrounding phase is required for meaningful volume compartmentalization and component enrichment to form functional assemblies (35, 39–42). In addition, a closer comparison between the condensate densities in *Fig. 2B* and *Fig. 2A* reveals that their differences primarily reside at columns of intermediate *J*_*p*_ values (*Fig. 2D* and *SIAppendix* Fig. S5E and F), indicative of different response patterns of surface condensates to *J*_*p*_ with and without obstacles.

The change in response of surface condensation to *J*_*p*_ upon obstacle introduction becomes clearer by examining horizontal rows of *Fig. 2A* and *B*. As shown in *Fig. 3A*, at fixed *J*_*m*_, without obstacles (see blue curve of *Fig. 3A*), surface condensation starts at *J*_*p*_ = 0.1 *k*_*B*_*T*, and the condensate density then increases gradually as *J*_*p*_ further increases. By contrast, with the introduction of 30% membrane obstacles, the onset of surface condensation is delayed to *J*_*p*_ = 0.35 *k*_*B*_*T*, and the condensate density varies more steeply to increasing *J*_*p*_, showing a sensitivity enhancement by a factor greater than three.

**Fig. 3.**
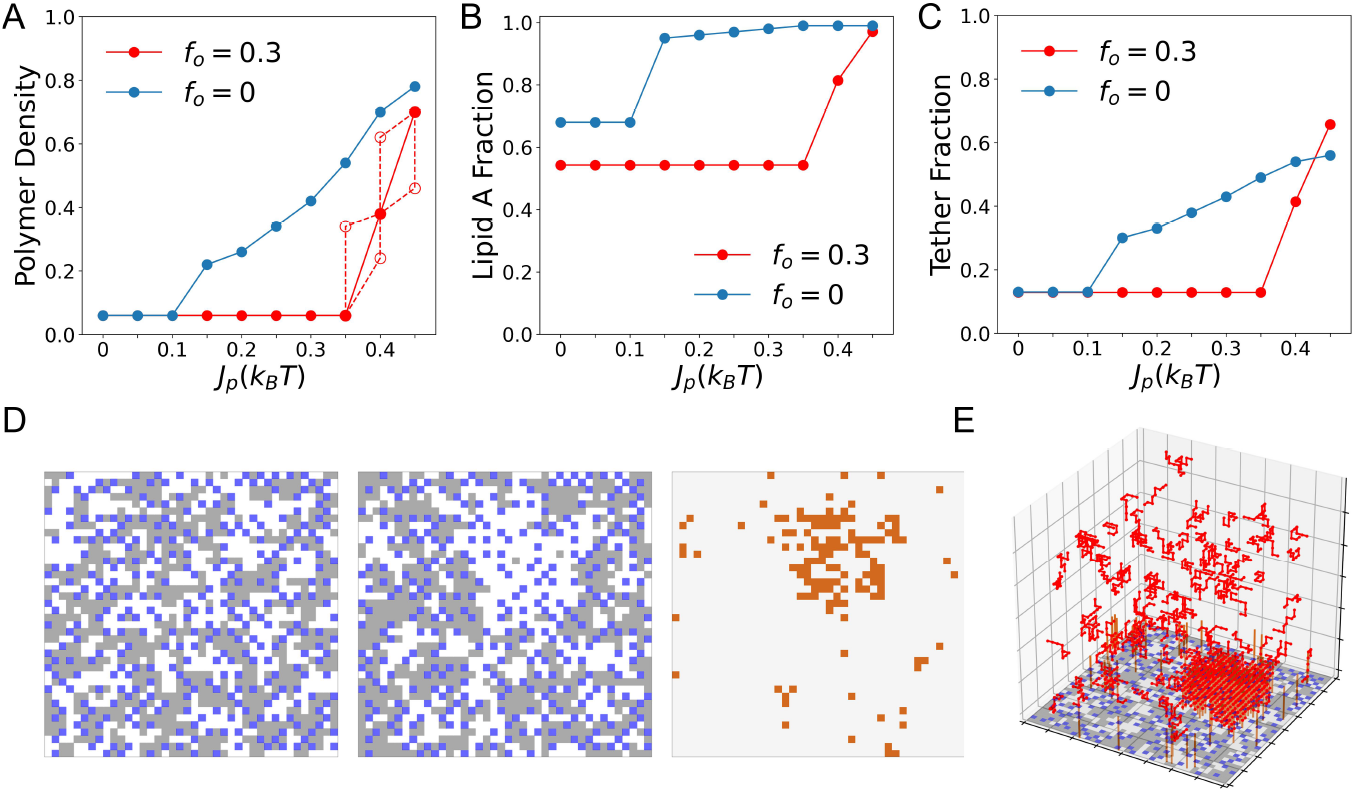
Membrane obstacles enhance the sensitivity and selectivity of surface condensation and its complementary membrane reorganization to *J*_*p*_; *f*_*o*_ represent the area fraction of membrane obstacles. (A) Polymer density of surface condensate as a function of *J*_*p*_ at fixed *J*_*m*_ = 0.55 *k*_*B*_*T* and *ρ*_*t*_ = 0.2. The empty red circles show data for *f*_*o*_ = 0.3 obtained with frozen membrane configurations. Specifically, the empty red circles at *J*_*p*_ = 0.4 *k*_*B*_*T* (lower one) and *J*_*p*_ = 0.45 *k*_*B*_*T* show condensate densities obtained when *J*_*p*_ is increased from 0.35 to 0.4 *k*_*B*_*T* and from 0.4 to 0.45 *k*_*B*_*T*, respectively, with membrane configurations taken from simulations with *J*_*p*_ = 0.35 and 0.4 *k*_*B*_*T*, respectively. Similarly, the empty red circles at *J*_*p*_ = 0.4 *k*_*B*_*T* (upper one) and *J*_*p*_ = 0.35 *k*_*B*_*T* show condensate densities obtained when *J*_*p*_ is set to 0.4 and 0.35 *k*_*B*_*T*, respectively, but with the membrane fixed at representative configuration selected from the trajectory with *J*_*p*_ = 0.45 and 0.4 *k*_*B*_*T*, respectively. (B) Fraction of type-A lipid (up spin) in all lipids beneath the surface polymer phases of (A). (C) Fraction of tethers on lipids beneath the surface polymer phases of (A). Data for *J*_*p*_ ⩽ 0.1 (0.35) *k*_*B*_*T* with *f*_*o*_ = 0 (0.3) describe the stable dilute surface polymer phase before the prewetting transition. (D) (Left) Snapshot of membrane simulation at *J*_*m*_ = 0.35 *k*_*B*_*T* with obstacle fraction of 0.2. (Middle) Snapshot of membrane in a simulation at *J*_*p*_ = 0.45 *k*_*B*_*T* and *J*_*m*_ = 0.55 *k*_*B*_*T* with obstacle fraction of 0.2. (Right) Corresponding tether distribution of the Middle snapshot. Chocolate and light grey squares represent positions with and without tethers. (E) Snapshot of the entire simulation system corresponding to the snapshot in D (Middle).

The presence of obstacles also contributes to regulating the accompanying membrane reorganization during surface condensation. As demonstrated in *Fig. 3B* and *C* (see also *SIAppendix*, Fig. S6), the presence of 30% obstacles enhances the sensitivity of the response of membrane composition beneath the surface condensate to *J*_*p*_, in terms of the concentrations of lipid A and tether, similar to that observed in *Fig. 3A* for condensate density. It should be noted that the realization of sensitive membrane reorganization beneath surface condensate by obstacles is functionally significant. Biological membranes have been proposed to be consist of domains of distinct composition and properties for accomplishing various cellular functions (23, 48–51). However, direct evidence for the existence of large lipid domains *in vivo* have been lacking, which was proposed to be due to their context-dependent nature and the multiplicity of their possible organizational states (52, 53). In line with this, our simulation shows that while the presence of macroscopic lipid domains is suppressed by obstacles, local lipid domain assembly is facilitated by a contacting surface condensate (demonstrated in *Fig. 3D* and *E*). Thus, the broad existence of protein obstacles in biological membranes may ensure that local lipid domains form only when needed.

The observed obstacle effects persist over obstacle types (see *SIAppendix*, Fig. S7), membrane components (see *SIAppendix*, Fig. S8D), and lipid environments (reflected by different *J*_*m*_ values, see *SIAppendix*, Fig. S8A-C). To better understand the physical origin of such obstacle effects, we reason that the obstacles can affect the membrane in two ways relevant to surface condensation. First, obstacles suppress membrane phase separation. Second, the presence of obstacles reduces the effective tether density on the entire membrane by a factor of (1-*f*_*o*_), especially in the lipid-mixed state of the membrane. Hence we set out to explore if we can reproduce the obstacle effects by repeating the obstacle-free simulations (blue curve of *Fig. 3A*) at reduced effective *J*_*m*_ and tether density (*ρ*_*t*_). From such analysis, we conclude that the sensitivity and selectivity enhancing effect of obstacles on surface condensation originates from creating membrane conditions unfavorable for forming dense polymer aggregates (see *SIAppendix* and Fig. S9 for detailed discussions).

### Mean-field Theory for the Obstacle Effects

To gain a deeper understanding of the obstacle effects on surface condensation, we then developed a theoretical model for our simulation system. Specifically, the free energy for the formation of surface condensation (*F*) is divided into three contributions: 1. aggregation of polymers from the bulk to the surface condensate; 2. reorganization of lipids and tether in the membrane beneath the surface condensate; and 3. the formation of interaction between tethers and the condensate polymers. We then minimize *F*, with and without obstacles, at different *J*_*p*_ values with respect to the density of the surface condensate (*ϕ*_0_) and the composition of the membrane underneath; these analyses enable us to dissect the effect of obstacles on the response of surface condensate and membrane composition to *J*_*p*_.

### Adapted Flory-Huggins Free Energy of Polymers Captures the Bulk Simulation Result

We describe the free energy of the polymers with an adapted Flory-Huggins theory (54, 55), in which the average free energy per monomer in the bulk (*f*_9_) is given by:

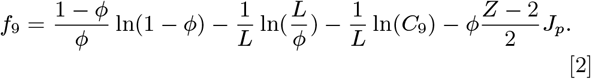

Here *ϕ* is the polymer concentration, *Z* = 6 is the cubic lattice coordination number, and *L* = 9 and *C*_9_ = 193, 983 are the length and the number of conformations with a given starting site of a single 9-monomer chain in the bulk (see *Methods* and *SIAppendix* for details). As plotted in *Fig. 4A*, at low *J*_*p*_ values, *f*_9_ increases monotonically with *ϕ*, indicating the dominance of entropy and a dilute solution. Yet, when *J*_*p*_ increases to ⩾ 0.55 *k*_*B*_*T, f*_9_(*ϕ*) becomes non-monotonic and develops a local minimum at a high *ϕ* value, which agrees with our bulk simulation result that phase separation starts at *J*_*p*_ = 0.55 *k*_*B*_*T* (see *SIAppendix*, Fig. S3). Thus, we take *f*_9_(*ϕ* = 0.08) = − 2.93 *k*_*B*_*T* as the free energy of a monomer in the bulk, which is equal to the value of *f*_9_(*ϕ*) for the local minimum at high *ϕ* value when *J*_*p*_ = 0.55 *k*_*B*_*T*. Accordingly, *ϕ* = 0.08 is taken as the bulk polymer concentration (*ϕ*^∞^). Considering that at low concentrations, the enthalpic contribution to the Flory-Huggins free energy is small, *ϕ*^∞^ = 0.08 and *f*_9_(*ϕ*^∞^) = − 2.93 *k*_*B*_*T* are considered as constants for *J*_*p*_ ⩽ 0.55 *k*_*B*_*T* (see *Fig. 4A*).

**Fig. 4.**
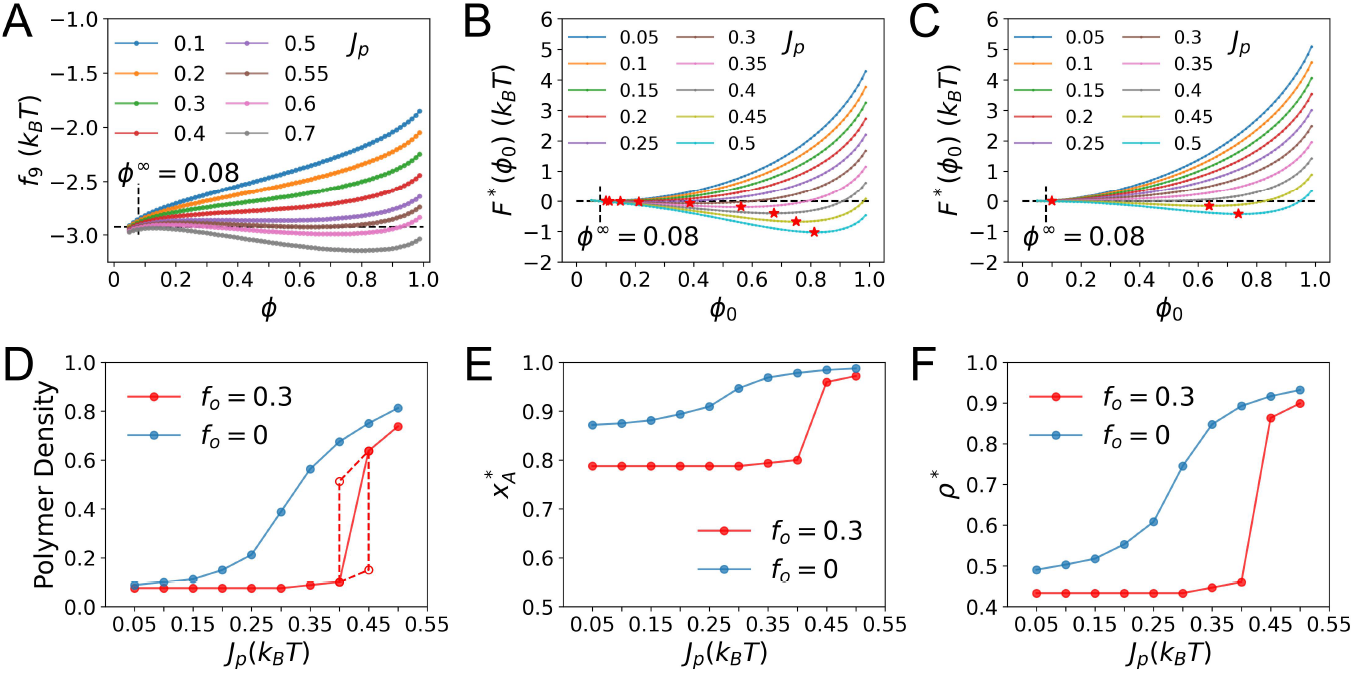
Mean-field theory (MFT) for the effect of obstacles on surface condensation. (A) Free energy per monomer as a function of polymer density in the bulk solution, *f*_9_ (*ϕ*), at different *J*_*p*_(*k*_*B*_*T*) calculated from the adapted Flory-Huggins theory. *ϕ*_∞_ represents the bulk polymer density. (B) Minimized *F* as a function of polymer density in the surface condensate *F*\*(*ϕ*_0_) at different *J*_*p*_(*k*_*B*_*T*) at *f*_*o*_ = 0. Red stars label the positions where *ϕ*_0_ minimizes *F*\*(*ϕ*_0_) at different *J*_*p*_(*k*_*B*_*T*), which have free energies lower than that of the dilute reference, leading to the formation of surface condensates. (C) Same as (B) but for *f*_*o*_ = 0.3. (D) Polymer densities; (E) lipid compositions; (F) tether concentrations of the surface condensates as a function of *J*_*p*_ at different obstacle fractions from the MFT calculations. In part (D) the left (right) empty red circle shows condensate density achieved in the calculation for *J*_*p*_ =0.4 (0.45) *k*_*B*_*T* while the membrane configuration is fixed at that corresponds to the minimum of *F*_*_(*ϕ*_0_) in a normal calculation at *J*_*p*_ =0.45 (0.4) *k*_*B*_*T*. Data in (D) to (F) at *J*_*p*_ < 0.1 (0.4) *k*_*B*_*T* when *f*_*o*_ = 0 (0.3) describe the dilute surface phase before the prewetting transition.

### Free Energy Analysis of Surface Condensation

The three contributions to *F* are divided into two parts *F* = *F*_3*D*_ + *F*_2*D*_, which are minimized separately like in the classical theory of wetting (18, 44, 45). Here *F*_3*D*_ is the free energy of forming a surface condensate of area *A* and density *ϕ*_0_ from a uniform dilute polymer solution at concentration *ϕ*^∞^. Specifically we have:

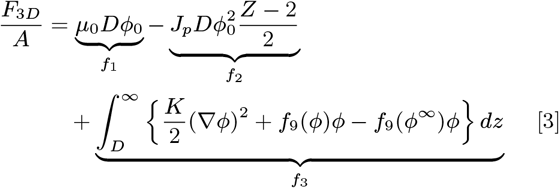

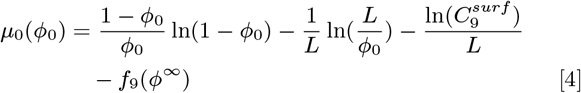

where *D* is the thickness of the surface condensate and 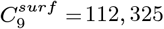 is the number of conformations of a 9-monomer chain with a given starting site confined in the surface condensate (see *SI Appendix* for details). *μ*_0_ in the term *f*_1_ accounts for the loss of conformational and translational entropies per monomer when bulk polymers are confined to the surface condensate. *f*_2_ is the enthalpic gain from the interaction between polymers in the surface condensate, and *f*_3_ is the interfacial energy between the bulk polymer solution and the surface condensate.

While *F*_3*D*_ describes what happens above the membrane, *F*_2*D*_ is the free energy of membrane reorganization and forming tether-polymer interaction during surface condensation. Specifically we have:

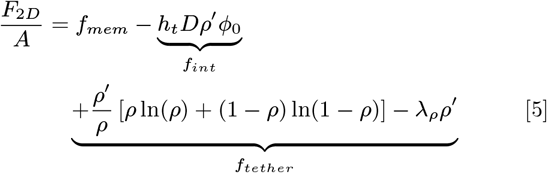

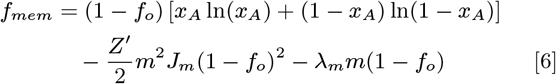

where 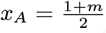 is the fraction of lipid A, *Z*^*′*^=4 the coordination number of 2D square lattice, *ρ* the local tether density on lipid A, 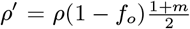 the overall tether density on the local membrane (regardless of lipid type), *h*_*t*_ the tetherpolymer interaction strength, *λ*_*ρ*_ the chemical potential of tether and *λ*_*m*_ is one half of the difference between the chemical potentials of lipid A and B. *f*_*mem*_ and *f*_*tether*_ represent the contribution to the membrane free energy from lipids and tether, respectively. *f*_*int*_ is the interaction energy between tethers and polymers (see *Methods* for derivations). The parameter values used in the MFT calculations are summarized in *SI Appendix* Table S1.

With *F*_3*D*_ and *F*_2*D*_ defined, we then minimize them independently at fixed *J*_*p*_ and *f*_*o*_ values for each *ϕ*_0_ to get 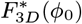 and 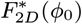. Therefore, the *ϕ*_0_ that minimizes their sum 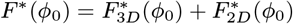, will be the density of the most stable surface polymer phase that forms at the given *J*_*p*_ and *f*_*o*_ (see *SI Appendix* and Fig. S10-11 for detailed discussions).

### Minimization of *F* (*ϕ*_0_)

Without obstacles, the sum of the minimized 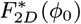 (black curve of *SI Appendix* Fig. S11A) and 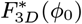 (*SI Appendix* Fig. S10) is plotted as *F*^*^(*ϕ*_0_) in *Fig. 4B*. At low *J*_*p*_ values, *ϕ*_0_ = *ϕ*^∞^ globally minimizes *F*^*^(*ϕ*_0_), indicating the absence of condensate formation. The situation of *ϕ*_0_ < *ϕ*^∞^ is ignored considering that we have a polymer-attracting membrane. When *J*_*p*_ reach 0.1 *k*_*B*_*T*, the local minimum of *F*^*^(*ϕ*_0_ = 0.1) (see the red star of the orange curve of *Fig. 4B*) becomes equal to *F*^*^(*ϕ*_0_ = *ϕ*^∞^), which marks the onset of surface condensate formation. As *J*_*p*_ further increases, the global minimum of *F*^*^(*ϕ*_0_) gradually shift to the right (see red stars in *Fig. 4B*), reflecting the gradual increase of condensate density as *J*_*p*_ increases.

On the other hand, when *f*_*o*_ is increased to 0.3, 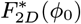 and its derivative adopt different values (*SI Appendix* Fig. S11C). Consequently, the formation of surface condensate now starts at a higher *J*_*p*_ (0.4 *k*_*B*_*T*), while featuring an increased density-to-*J*_*p*_ sensitivity, as manifested by the red star locations on the *F*^*^(*ϕ*_0_) curves in *Fig. 4C*. These observed differences in the condensate-to-*J*_*p*_ response, as well as the membrane composition-to-*J*_*p*_ response from adding obstacles are further summarized in *Fig. 4D* − *F*, which well recapitulate the corresponding observations from the simulation results (see *Fig. 3A* − *C*).

### Physical Origin of the Obstacle Effects

The formation of surface condensation consists of several contributions: 1. the confinement of polymers from a dilute bulk solution into the surface condensate, which is unfavorable due to the loss of conformational and translational entropies of the confined polymers and the distortion of polymer concentration profile above the condensate; 2. formation of favorable polymer-polymer interactions in the surface condensate, where the polymer concentration is much higher than in the bulk solution; 3. concentration of specific lipids and tethers to the membrane beneath the condensate, which is unfavorable due to the loss of their translational entropies, especially above the critical temperature of the membrane; 4. formation of favorable tether-polymer interactions in the surface condensate. As such, surface condensate emerges only when the free energy gain of contributions 2 and 4 outweighs the loss of contributions 1 and 3. Therefore, to achieve a surface condensate of a given density *ϕ*_0_ and certain membrane composition beneath it, which involve fixed free energies of contributions 1 and 4, the enthalpic gain of contribution 2 needs to overcome the entropic loss of contribution 3. The presence of obstacles, which suppresses the concentration of membrane components in contribution 3 (as manifested in the simulation results and Eq. **5**-**6**), thus requires stronger polymer-polymer interactions to drive the formation of the surface condensate. This explains the right shift of the *ϕ*_0_ − *J*_*p*_, *x*_*A*_ − *J*_*p*_ and *ρ* − *J*_*p*_ curves in *Fig. 3A* − *C* and *Fig. 4D* − *F*, or the enhancement of selectivity towards *J*_*p*_ due to membrane obstacles for the formation of surface condensates.

While the above argument elucidates the delay of the onset of surface condensation to higher *J*_*p*_, it doesn’t guarantee the enhancement of sensitivity. In fact, now that it’s harder to drive the membrane reorganization by increasing *J*_*p*_ in the presence of obstacles, one might expect it is even harder to drive fast condensate assembly. To gain further insights, we reason that instead of only focusing on how membrane responds to *J*_*p*_, we should also inspect how the change of local membrane composition influences the condensate. Noticeably, in both the simulation and the MFT results, rapid condensate density increase and membrane reorganization are observed simultaneously (*Fig. 3A* − *C* and *Fig. 4D* − *F*). Without obstacles, when the membrane is close to phase separation on its own, a modest enrichment of polymers in the surface condensate at low *J*_*p*_ could already effectively induce its reorganization. The membrane composition then responds to *J*_*p*_ or condensate density in a gradual manner (see blue curves in *Fig. 3A* − *C* and *Fig. 4D* − *F*). By contrast, when the obstacles suppress lipid phase separation and scale down the effective lipid A-polymer affinity (see Eq. **5**), membrane reorganization only starts at a stronger *J*_*p*_ in a shallow manner and then is driven progressively higher (see red curves in *Fig. 3A* − *C* and *Fig. 4D* − *F*). This leads to the realization that, with obstacles, condensate density and membrane composition become more sensitive to increasing *J*_*p*_ because they increase co-operatively. More specifically, it is exactly because it’s harder to drive membrane reorganization due to obstacles that membrane composition responds to the increase of *ϕ*_0_ (*J*_*p*_) in a co-operative manner (Eq. **5**); i.e., the positive co-operativity between the coupled growth of condensate and local membrane domain ultimately leads to the sensitivity enhancement. This co-operative mechanism is further demonstrated by the red dashed lines in *Fig. 3A* and *Fig. 4D*, where when *J*_*p*_ increases with frozen membrane configuration, condensate density responds much less, and vice versa.

### General Significance of Membrane-surface polymer Co-operativity to Surface Condensation

The principle of positive co-operativity is quite general. For any particular polymer states, e.g., characterized by the range of *J*_*p*_ values, other parameters can be tuned collectively to modulate the degree of this positive co-operativity. In this section, we investigate several other ways of tuning membrane-surface polymer co-operativity to verify the generality of the principle. As the problem is ultimately concerned with two-body co-operativity, the model membranes studied in this section consist of only lipids A and B (without obstacles or tethers), with a direct attraction between lipid A and surface polymers. We then adjust the membrane-surface polymer co-operativity in four distinct ways by varying: 1. lipid A fraction *f*_*A*_; 2. lipid-lipid interaction strength *J*_*m*_; 3. strength *h*_*t*_ and 4. range *l* of lipid A-polymer attraction to investigate how they affect condensate property (see *Methods* for details).

The four ways of tuning membrane-surface polymer co-operativity could be further classified into two categories based on their effects: 1. varying *f*_*A*_ and *J*_*m*_ changes the propensity of the membrane to phase separate by itself (see *SI Appendix* Fig. S12); 2. varying *h*_*t*_ and *l* changes the ability of surface polymer to reorganize the underneath membrane (see Eq. **15** and **4**) and *SI Appendix* for more discussions). Three regimes of surface condensation are observed depending on the state of the membrane and polymer in terms of these two aspects.

As summarized in Fig. 5 and *SI Appendix* Fig. S13 and S14, when the membrane is too far from phase separation, prewetting is suppressed and a dilute surface polymer phase dominates at all *J*_*p*_ values (see black lines of 5*A* − *D* and *SI Appendix* Fig. S13 A-D). On the other hand, when the membrane is close to phase separation, a dilute surface polymer phase at low *J*_*p*_ value is sufficient to induce local membrane reorganization and the prewetting transition (see blue curves of 5*A* − *D* and *SI Appendix* Fig. S13 A-D). The condensate density responds in a gradual manner to varying *J*_*p*_, with the membrane composition beneath the condensate remaining largely constant.

**Fig. 5.**
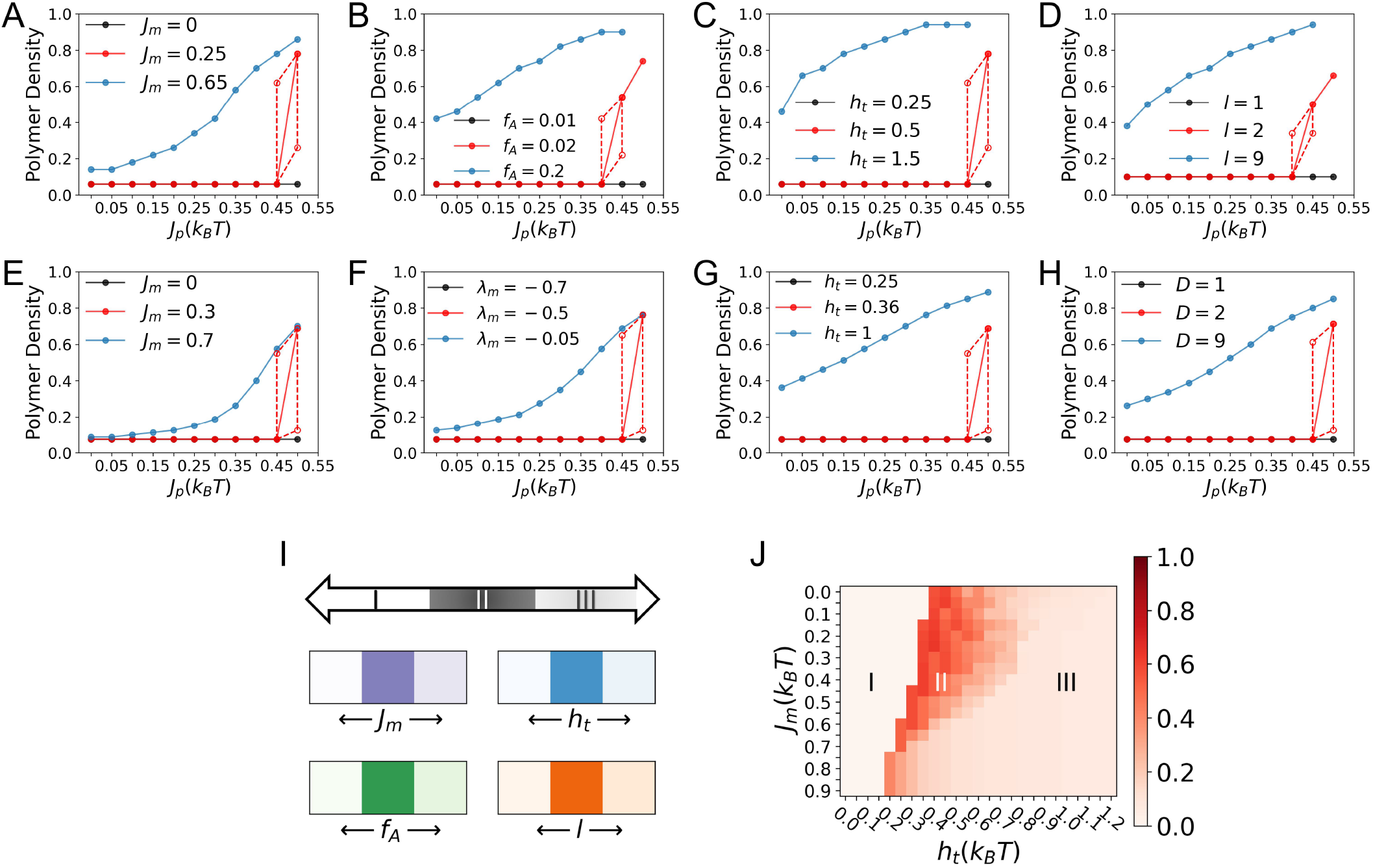
The connection between membrane-surface polymer co-operativity and surface condensate property is generally applicable to other model membrane-polymer systems (see *Methods* for details) as revealed by Grand Canonical Monte Carlo (GCMC) simulations and a mean-field theory (MFT). GCMC results for the surface condensate density as a function of *J*_*p*_ (A) at three *J*_*m*_ values; (B) at three *f*_*A*_ values; (C) at three *h*_*t*_ values; (D) at three *l* values; with other parameters fixed (see *SI Appendix* Table S2). MFT results for surface condensate density as a function of *J*_*p*_ (E) at three *J*_*m*_ values; (F) at three *λ*_*m*_ values; (G) at three *h*_*t*_ values; (H) at three *D* values with other parameters fixed (see *SI Appendix* Table S3). The black data points in (A)-(H) describe the stable dilute surface polymer phases before the prewetting transition. The empty red circle data in (A)-(D)/(E)-(H) are similarly defined as those in Fig. 3*A*/Fig. 4*D*. The lower empty red circle of (E)-(H) (upper empty red circle of (F)) is shifted up (down) by 0.05 to avoid overlapping for better visualization. (I) A schematic illustration of the 1D phase diagram that summarizes the three regimes of surface condensation, which depends on the degree of membrane-surface polymer co-operativity. The lower panels show that a system can be shifted among the three regimes by varying different parameters. The darkness of the colors of the lower four panels from left to right reflects the highest *ϕ*_0_ change over a *J*_*p*_ change of 0.05 *k*_*B*_*T* of the black, red and blue curves of (A)-(D). (J) A two-dimensional phase diagram obtained from MFT calculations spanned by *J*_*m*_ and *h*_*t*_. The darkness of the color reflects the highest condensate density change over a *J*_*p*_ change of 0.05 *k*_*B*_*T* with *λ*_*m*_ = −0.24, *D* = 5.

When parameters are tuned, however, so that membrane reorganization is not favored yet a prewetting transition is still possible, surface condensation only starts at a high *J*_*p*_ value. Under such situation, the condensate density and the local lipid domain grow co-operatively as *J*_*p*_ further increases, and the coupled growth leads to their sensitive response to *J*_*p*_ (see red curves of 5*A* − *D* and *SI Appendix* Fig. S13 A-D). Results from the free energy analysis of the mean-field theory agrees well with the simulation results, as summarized in 5*E* − *H, SI Appendix* Fig. S13 E-H and Fig. S14 (see Methods and *SI Appendix* for more discussions).

These results confirm that the connection between membrane-surface polymer co-operativity and surface condensate property is not limited to the discussion of membrane obstacle effect, but more general. Indeed, the observations here suggest that a membrane-polymer system can be classified into one of the three regimes based on the degree of membrane-surface polymer co-operativity as summarized in *Fig. 5I* : In regime I, the membrane is far away from phase separation that surface condensation is suppressed; in regime III, the membrane is close to phase separation that leads to surface condensation of low selectivity and sensitivity; in the intermediate regime II, surface condensation of high selectivity and sensitivity towards polymer property is observed. As summarized in the lower panels of *Fig. 5I*, the key factor of membrane-surface polymer co-operativity can be tuned in various ways, thus leading to multiple mechanisms through which the surface condensate formation and property can be regulated.

Our analyses highlight that for the discussion of surface condensate regulation, membrane’s tendency to phase separate should be considered together with the polymer’s ability to reorganize the membrane (see *Fig. 5A* − *H* and *SI Appendix* Fig. S13 and S14). This is explicitly illustrated in *Fig. 5J*, where *J*_*m*_ and *h*_*t*_ together determine the boundary that separates the three regimes of surface condensation in this two-dimensional parameter space (see *SI Appendix* for more discussions). More broadly, it is the collective effect of all relevant parameters of the system that determines the degree of membrane-surface polymer co-operativity, which ultimately regulates the formation and property (e.g., density) of surface condensate (see also *SI Appendix* Fig. S15-17).

Finally, although the current study focuses on biomolecular system, the general principle that positive co-operativity between the coupled growth of two order parameters results in their faster increases applies to broader contexts. For example, the positive reciprocal effects between incidental news exposure via social media and political participation observed in communication studies reflects similar principles only with polymers and lipids replaced by social media exposure and political participation (56), respectively. Such a general principle is also demonstrated with an intuitive example of tennis practice (see *SI Appendix* and Fig. S18).

### Concluding Remarks

In recent years, biological membranes have been recognized to play a regulatory role in the formation of biomolecular condensates (13–17), especially in the context of cell signaling. The general physical principles and molecular details that govern the robustness and sensitivity of such regulations, however, remain to be elucidated. For example, the recent study of Machta and co-workers (18), which played a major role in inspiring the current work, highlighted the potential significance of the membrane being close to its critical point (57); it was shown that, as the membrane approaches its critical point, the range of polymer interaction strength that leads to pre-wetting transition is greatly expanded. While this was an interesting observation, the insensitivity of the pre-wetting transition to the polymer interaction strength implies the lack of selectivity, making the mechanism less than ideal from a functional perspective. Moreover, as discussed here and in previous work (32, 33), realistic cellular membranes, which are rich in proteins, are unlikely to undergo macroscopic phase separation under ambient conditions. Additionally, while previous studies mostly focused on the conditions that favor surface condensation, the regulatory mechanism of condensate property (e.g., density), which affect the functions of MLOs, remains relatively unexplored (18, 19, 35–45).

Motivated by these considerations, in this work, we have first studied the effect of protein obstacles on the coupled phase behaviors of biological membranes and surface biopolymers through GCMC simulations and a mean-field theory. We confirmed previous theoretical analysis (32, 33) that the presence of protein obstacles, especially immobile ones, suppresses phase separation of lipid membrane. Despite deviation from the critical point, the local membrane composition responds co-operatively to the condensation of surface polymers, which is in turn further promoted by the local enrichment of specific lipids; co-operativity in the coupled growth thus leads to enhanced sensitivity and selectivity of local membrane reorganization and pre-wetting transition to the properties of the polymer, represented by the interaction strength *J*_*p*_ in the current model. The key role of the obstacles in enhancing the sensitivity and selectivity is, in fact, to push the membrane away from conditions that strongly favor surface condensation, leaving opportunities for polymer properties to contribute.

This mechanism is further tested with several other membrane-polymer systems, which help establish the general principle that relates the degree of membrane-surface polymer co-operativity and condensate property regulation. As far as parameters of the system are tuned to generate positive co-operativity in the coupled growth of local membrane domain and surface condensate, high sensitivity and selectivity of condensate regulation is realized.

The sensitivity and selectivity of the surface condensate to the polymer properties are functionally relevant; the polymer properties can be modified by variations in sequence, PTM state or local environmental variables such as pH. For example, a high level sensitivity of phase separation to these changes is essential for cells to produce digitized output in processing cytoplasmic or external signals through the mechanism of surface condensation (58). Previous studies have revealed that the assembly of condensates of the linker for activation of T cells (LAT) in the T cell receptor (TCR) signaling pathway responds nonlinearly to the phosphorylation states of its tyrosine sites; this feature was proposed to be correlated with the selectivity and sensitivity of TCR antigen discrimination (16, 17, 59, 60). The pH sensitivity of the condensate formation by the prion protein Sup35 was found to promote yeast cell fitness (61). Methylation of arginine sites was suggested to be an effective physiological regulator of fused in sarcoma (FUS) phase behavior (62); the density of FUS condensate influences its propensity for fibrillization, which is linked to neurodegenrative diseases (36–38). The condensation state of epidermal growth factor receptor (EGFR) and the adaptor protein Grb2 was revealed to be sensitive to their phosphorylation states, which could regulate downstream signal propagation to the mitogen-activated protein kinase (MAPK) pathway (63), and phosphorylation is also believed to regulate gephyrin mediated clustering of receptors in inhibitory synapses via charge-charge interaction driven phase separation (64). Our analyses suggest that membrane obstacles or other ways of enhancing co-operativity may contribute constructively to the sensitivity and selectivity of signal transduction processes mediated by surface phase separation.

Our observation that surface condensate formation helps promote local lipid segregation agrees with the recent experimental observation that aggregation of attached surface biomolecules can drive the formation of phase separated lipid domains in membranes at temperature well above its *T*_*c*_ (65–69) Although our model system is minimal and surface condensation in biology may feature much greater complexity, the results from the current study highlights that creating positive co-operativity between membrane and surface phase behaviors represents a general strategy for enhancing the sensitivity and selectivity of signal transductions. We anticipate that such predictions can be experimentally tested *in vitro* through reconstituting phase separating polymer systems in the presence of multi-component lipid membranes with tunable lipid and obstacle compositions, range of lipid-polymer attractions (e.g., modifying range of electrostatic interactions through changing salt concentration), and strength of lipid-lipid or lipid-polymer interactions (e.g., through post-translational modifications) (13).

The obstacles explored here are inert in that they do not feature any preferential interactions with any lipid or polymer. In reality, protein obstacles may have specific features that further modulate their impact on the coupled membrane/biopolymer phase behaviors. For example, obstacles with preferable interaction with selected lipids, which has been shown to eliminate membrane phase separation at a rather low density, could potentially be an effective way for implementing location selectivity in surface condensate formation (33, 34). In addition, obstacle’s structural properties can also affect surface condensation formation. For example, the protruding parts of obstacles or protein-induced curvatures may introduce surface roughness that places a length-scale threshold on proteins that are capable of conformal coating and the subsequent condensation (70–72). Finally, how non-equilibrium processes, which are prevalent in signal transductions, contribute to the coupled phase behaviors of complex membrane and surface biopolymers and, therefore, sensitivity and selectivity of cellular responses is a fascinating topic for future explorations.

## Methods

### MC Simulations of the Membrane

The lipid membrane is modeled by fixed composition 2D Ising model depicted in the main text with Hamiltonian given in Eq. **1** repeated below.

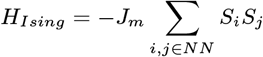

Simulation for each membrane condition (*J*_*m*_ and the concentration of immobile/floating obstacles) consists of 10^6^ MC sweeps through all lipid and mobile obstacle sites. MC moves are accepted with the Metropolis probability 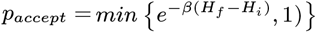, where *H*_*i*_*/H*_*f*_ is the energy of the system before and after the proposed move and 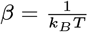 of the membrane is averaged over configurations generated every 100 MC sweeps.

### GCMC Simulations of the Polymer-Membrane System

The coupled polymer-membrane system is modeled as depicted in the main text, whose Hamiltonian is given by Eq. **7** below.

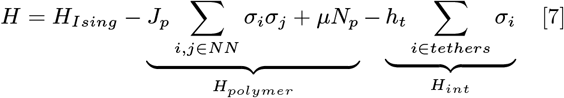

Here, *σ*_*i*_ = 1/0 if a lattice site *i* above the Ising membrane is occupied/unoccupied by a monomer. NN interaction between two connected monomers from the same polymer chain is not counted. A lattice site can not be occupied by more than one monomer. *μ* = −19*k*_*B*_*T* is the chemical potential of a single 9-monomer chain and *N*_*p*_ is the total number of polymer chains in the simulation box. *h*_*t*_ is the interaction energy between polymers and tethers. *i tethers* include all lattice site *i* that is occupied a tether, thus a polymer-tether interaction forms when a monomer occupies the same lattice site as a tether. A tether must be attached to a lipid-A membrane site and cannot overlap with other tethers. The simulation box contains 40 × 40 × 30 (*xyz*) lattice sites and has PBC in the *x* − *y* plane with non-penetrable boundaries in the *z* direction.

In the simulation of the coupled system, each MC sweep is divided into two sequential steps. The first step contains 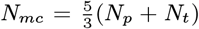 MC moves, where *N*_*p*_ is the number of polymer chains in the system before the start of the MC sweep, and *N*_*t*_ is the total number of tethers in the system (fixed at initialization). 40% of the *N*_*mc*_ MC moves are assigned to the addition and deletion moves of polymers equally, while the remaining 60% are assigned to polymer moves and tether moves in the ratio of *N*_*p*_ : *N*_*t*_. All *N*_*mc*_ moves are then carried out in a randomized sequence. The second step of the MC sweep is the membrane simulation, which sweeps through all lipid and mobile obstacle sites like described in the previous section, except that tether attached lipid-A sites are not allowed to move.

In a polymer addition move, a 9-monomer chain (9-mer) is selected randomly from a pre-built pool that contains all possible 9-mer configurations. This selected 9-mer is then proposed to be added to the simulation box with its center to be placed at a randomly selected lattice site. The probabili of acceptance is 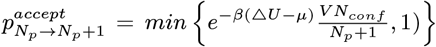 where Δ*U* is the energy change of the system by the proposed move, *N*_*p*_ the number of 9-mers in the system before the move, V the system size and *N*_*conf*_ the number of all possible 9-mer configurations (73).

In a polymer deletion move, a randomly selected 9-mer from the current simulation box is proposed to be deleted with 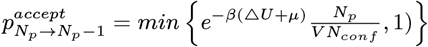. Addition or dele-tion of a 9-mer is immediately rejected if it leads to monomer overlap, change of the number of inter-chain interactions or polymer-tether interactions in the system. Special care is taken to treat the boundaries in the *z* direction in terms of the values of *N*_*conf*_.

In a tether move, a randomly selected tether is proposed to translate one lattice site on the *x* − *y* plane with the Metropolis acceptance ratio. A tether can only be moved to neighboring lipid-A sites.

In a polymer move, for the majority of the time, a randomly selected 9-mer is proposed to do reptation move, in which a bond is removed from one end of the chain and glued to the other end at a random direction. A reptation move is accepted with the Metropolis probability while no monomer overlap is allowed. With a probability of 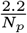 we propose cluster move of the selected 9-mer. A cluster is identified as the collection of all polymers connected to the selected 9-mer directly or indirectly through monomer-monomer NN contact. The cluster is proposed to move one lattice site at a random direction with the Metropolis acceptance ratio, while moves that result in the growth of the cluster are rejected to satisfy detailed balance (46).

In simulations of systems without tethers, a direct attraction between lipid A and polymer is implemented. Effectively, it can be thought of as every lipid A has a tether of length *l* for polymer attraction. All MC moves then stay the same as stated above besides the absence of tether moves.

Surface condensate density is obtained through analyzing polymer density distribution of 5 × 5 × 5 (*xyz*) surface regions (*P* (*ϕ*_0_)) averaged over configurations generated every 100 MC sweeps in a typical simulation of 10^6^ MC sweeps. As demonstrated in *SI Appendix* Fig. S19, in a system without surface condensation, *P* (*ϕ*_0_) is unimodally distributed, indicating the dominance of a dilute surface phase. When conditions of the system is tuned to enable surface condensation, *P* (*ϕ*_0_) becomes bimodally distributed with two peaks at different *ϕ*_0_ values, indicating phase separation. The condensate density is then identified as the higher *ϕ*_0_ at the two *P* (*ϕ*_0_) peaks (see *SI Appendix* Fig. S19). In simulations with *l* = 2, 3, 7 and 9, *P* (*ϕ*_0_) analysis described above is done for 5 × 5 × *z* surface regions with *z* = 3, 3, 7 and 9 as the condensate thickness changes with *l* (see *SI Appendix*).

The effectiveness of such simulation scheme in exchanging polymers between the dense condensate and the dilute solution is further demonstrated in *SI Appendix* Fig. S20

### Adapted Flory-Huggins Theory

In the classical Flory-Huggins theory, the number of independent configurations for arranging *N* polymers of length *L* in *M* lattice sites is given by:

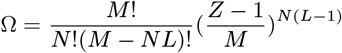

where *Z* is the coordination number of the lattice. In this work, we replace the part for counting conformational entropy (*z* − 1)^*N*(*L*−1)^ with 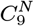 to avoid intra-chain overlap. Combined with the mean field energy 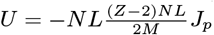 we have the free energy per monomer given in Eq. **2**:

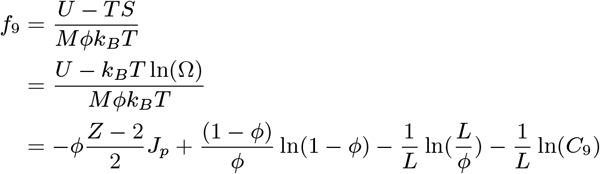

where *ϕ* = *NL/M* is the polymer density.

### Minimization of *F*_3*D*_ in Mean-field Theory

At given *ϕ*_0_ and *J*_*p*_ (*f*_1_ and *f*_2_ are fixed), the minimization of *F*_3*D*_ is just the minimization of the integration in the term *f*_3_ over the polymer density profile *ϕ*(*z*) above the surface condensate. To satisfy 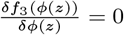 we have:

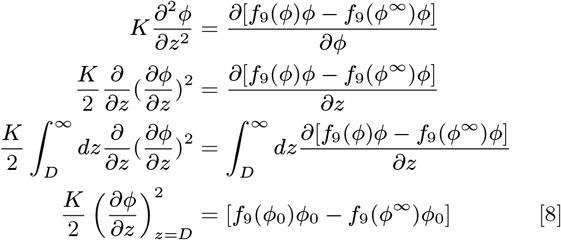

Substituting Eq. **8** into *f*_3_ we have:

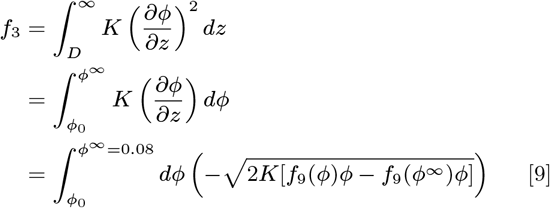

Eq. **9** (minimized *f*_3_ for a given *ϕ*_0_ and *J*_*p*_) can then be easily evaluated numerically.

### Lipid membrane Free Energy

The lipid free energy is obtained through a mean-field treatment. Specifically, the energy and entropy of a membrane of area *A* (lattice sites), with obstacle fraction *f*_*o*_ and lipid A fraction 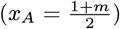 is given by

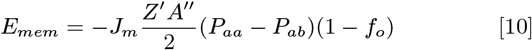

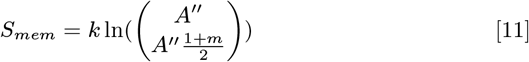

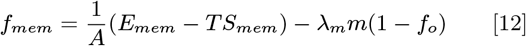

where *A*^*′′*^ = *A*(1 − *f*_*o*_) is the number of total lipids, 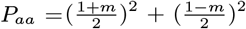 and 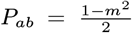 are the probability of a bond between like and unlike lipid pairs. With Stirling approximation, it can be shown that *f*_*mem*_ of Eq. **12** is equal to *f*_*mem*_ in Eq. **6**. *f*_*mem*_ derived here leads to the same critical point 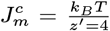 for 2D Ising model as usually seen in the MFT (see *SI Appendix* Fig. S21) (74).

### Tether Free Energy

The entropy of having a fraction *ρ* of the *A*^*′*^ lipid sites to be attached to a tether (non-interacting) is:

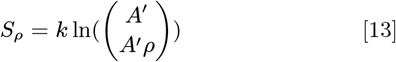

where 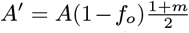 is the number of up spin sites within area *A*. Then the free energy contribution per unit area from tether arrangements is:

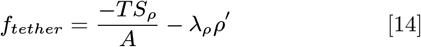

With Stirling approximation, it can be shown that *f*_*tether*_ of Eq. **14** is equal to *f*_*tether*_ in Eq. **5**.

### MFT of Tether-Free Systems

The above depicted MFT can be easily adjusted to describe the free energy of tether-free systems, by setting *f*_*o*_ = 0 and deleting contributions from tethers. Specifically, the expression of *F*_3*D*_ stays the same as in Eq. **3** and **4**, while the value of 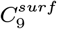 changes when *l* changes, and is evaluated in the same way as *l* = 5 (see *SI Appendix* for details). The modified expression of *F*_2*D*_ is given below.

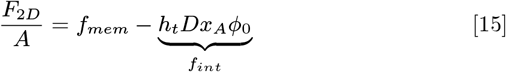

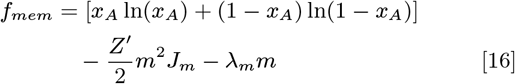

## Data Availability

The code used for GCMC simulations and MFT calculations in this study is available at https://github.com/liuzhbu/Co-operative-surfacecondensation. All data is included in the manuscript and/or supporting information.

## ACKNOWLEDGMENTS

The work is supported in part by grants NSF-DMS1661900 and NSF-CHE-2154804 to QC, and AY acknowledges support from grant NSF-CHE-1856595. Computations were conducted on the Shared Computing Cluster, which is administered by Boston University’s Research Computing Services (URL: www.bu.edu/tech/support/research/).

