## Supporting Information for "Sensitive and Selective Polymer Condensation at Membrane Surface Driven by Positive Co-operativity"

**This PDF file includes:**

Supplementary text
Figs. S1 to S21
Tables S1 to S3
SI References

### Supporting Information Text

### Effect of protein obstacles on membrane phase separation

The phase behavior of the membrane lipids is studied using the probability distribution function $P(x_A)$, of local molar fractions of lipid A ($x_A$) in regions of $5 \times 5$ lattice sites. As shown in Fig. S1B, without obstacles, $P(x_A)$ peaks around $x_A = 0.5$ at low $J_m$, indicating a well-mixed state of the lipids driven by entropy. As $J_m$ increases, the peak at $x_A = 0.5$ gradually broadens, until it goes beyond a critical value of $J_m^c = 0.35\ k_BT$ (green curve in Fig. S1B), when enthalpic effect dominates and $P(x_A)$ becomes bimodally distributed at two extreme values of $x_A$, reflecting the coexistence of two types of phase separated regions enriched in either lipid A or B. At different $J_m$ values, $P(x_A)$ remains symmetric about $x_A = 0.5$ as it should due to the spin-flip symmetry of the Hamiltonian. When obstacles are introduced, the qualitative characteristics of the $P(x_A)$ curves described above remain unchanged. Yet, as the fraction of obstacles increases from 0 to 0.1, 0.2 and 0.3, the critical membrane coupling $J_m^c$ increases from 0.35 $k_BT$ to 0.4, 0.5 and 0.7 $k_BT$, respectively (Fig. S1C-E). This can be understood from the fact that with the introduction of obstacles, the effective coordination number of the square lattice decreases, which is directly proportional to the critical temperature ($\propto \frac{1}{J_m^c}$) of Ising model in the mean-field theory (MFT)(1). Moreover, at high obstacle densities, the suppression of critical temperature ($T_c$) is much more dramatic than predicted by MFT (2). Intuitively, in an extreme case where the lipids are segregated into stripes by inert obstacle boundaries, the system becomes effectively one-dimensional (1D), which has no long-range order or a finite critical point(1).

The suppression of phase separation by floating obstacles is far less prominent than that from fixed obstacles (see Fig. S2). This can be explained by realizing that floating obstacles exert more flexible constraints on the reorganization of lipids compared with fixed obstacles, and thus are less effective at preventing their phase separation as $J_m$ increases . All conclusions drawn from analyzing $P(x_A)$ of regions of $5 \times 5$ lattice sites remain the same in the analysis of $P(x_A)$ for $10 \times 10$ regions, except a significant increase of $J_m^c$ in all situations, reflecting that it's harder to form larger phase separated domains in a fixed composition membrane or that correlation length increases with $J_m$ (see Fig. S3).

### Effect of protein obstacles on surface condensation originates from modifying membrane properties

As seen in Fig. S1B and S1E, the $P(x_A)$ of a membrane with 30 percent obstacles at $J_m = 0.55\ k_BT$ recapitulates that of a membrane without obstacles at $J_m = 0.3\ k_BT$. However, reducing $J_m$ from 0.55 $k_BT$ to 0.3 $k_BT$ in an obstacle-free membrane system has a much smaller effect on surface condensation than adding 30 percent obstacles (see green curves of Fig. S9). When we further reduce $\rho_t$ by a factor of (1-0.3) on top of the $J_m$ reduction, their combined effect becomes close to that of 30 percent obstacles (see purple curves of Fig. S9). The remaining slight difference between the purple and red curves of Fig. S9 can be attributed to the hard limit of local membrane composition placed by obstacles, which can not be achieved by modifying $J_m$ or $\rho_t$. Such analysis shows that the sensitivity and selectivity enhancing effect of obstacles on surface condensation originates from creating membrane conditions unfavorable for forming dense polymer aggregates. To verify this observation, we further performed obstacle-free simulations (blue curve of maintext Fig.3$A$) at an increased tether density, which indeed resulted in a broader and slower response of condensate density to $J_p$ (see pink curves of Fig. S9).

In line with the discussions above, floating obstacles also enhance the sensitivity and selectivity of surface condensation (Fig. S7) because they suppress membrane phase separation in a similar way as fixed obstacles do (see the previous section). On the other hand, as floating obstacles have a less prominent effect on membrane phase behavior, their effect on surface condensation is accordingly smaller (see main text Fig.2$B$, Fig. S5A-C and Fig. S7A-D) compared with that of fixed obstacles at the same concentration. However, the effect of fixed obstacles can be reproduced by floating obstacles of higher concentration. As demonstrated in Fig. S7E, 50 percent floating obstacles well recapitulate the effect of 30 percent fixed obstacles.

From a broader perspective, although the effects of fixed and floating membrane obstacles on surface condensation are subject to the restrictions set in this study (inert obstacles and membrane of critical composition), they serve as two unique ways of tuning membrane-surface polymer co-operativity, in parallel to the other variations we've explored (see the main text section "*General Significance of Membrane-surface polymer Co-operativity to Surface Condensation*"), and the demonstrated "obstacle effects" are in line with the general principle connecting the degree of membrane-surface polymer co-operativity and surface condensate regulation established in the current study. Moreover, as proven here and in previous works, (2–4), realistic cellular membranes, which are rich in proteins, are unlikely to undergo macroscopic phase separation under ambient conditions. Thus it is important to consider a broad range of membrane conditions when analyzing membrane mediated condensate formation.

### Evaluation of $C_9$ and $C_9^{surf}$

**$C_9$.** The number of all possible conformations of a 9-monomer chain (9-mer) in the bulk ($C_9 = 193983$) is evaluated numerically. Specifically, the central monomer of a 9-monomer chain is placed at the center of a large simulation box, and then $C_9$ is evaluated as the number of distinct ways to arrange the rest 8 monomers without self-overlapping.

**$C_9^{surf}$.** The number of all possible conformations of a 9-monomer chain in the surface condensate ($C_9^{surf} = 112325$) is evaluated numerically. As demonstrated in Fig. S4E, the thickness of the surface condensate in the prewetting regime is five lattice sites (which is the same as the length of tethers). Thus, $C_9^{surf}$ is evaluated as:

$$C_9^{surf} = \frac{\sum_{i=1}^{5} C^i}{5}$$

where $C^i$ is defined as the number of all possible 9-mer conformations whose central monomer is placed at a distance of i lattice sites from the membrane (z=i) without self-overlapping or penetration of planes at z=1 and z=5.

In the GCMC simulations of tether-free systems, when the range of lipid A-polymer affinity $l$ changes, the thickness of the surface condensate changes accordingly (see Fig. S17E). To model such effect in the corresponding MFT analysis (maintext Fig. 5$H$, Fig. S13H and Fig. S14D, S14H and S14K), the surface polymer phase thickness $D$ is set to 1, 2 and 9 for modeling $l = 1$, 2 and 9. The $C_9^{surf}$ value in these three cases are evaluated to be 2958, 30104 and 148563 using the same method described in the previous paragraph.

### Minimization of $F_{3D}(\phi_0)$

As shown in maintext Eq. **3** and Eq. **4**, when $J_p$ and $\phi_0$ are defined, the values of $f_1$ and $f_2$ are then fixed, and thus the minimization of $F_{3D}$ involves only the minimization of $f_3$ over the polymer density profile $\phi(z)$ above the surface condensate (see maintext *Methods* for detail). The minimized $F^*_{3D}(\phi_0)$ at different $J_p$ values are plotted in Fig. S10, which are all positive definite functions that increase monotonically with $\phi_0$. This is because without bulk phase separation, $\phi^\infty$ minimizes $f_9$ for $\phi \geqslant \phi^\infty$, and so the more polymers that are confined to the surface phase, the higher the free energy penalty. Nonetheless, $F^*_{3D}(\phi_0)$ becomes less positive when $J_p$ increases, due to the increased enthalpic gain from polymer-polymer interaction in polymer dense regions.

### Minimization of $F_{2D}(\phi_0)$

As seen in maintext Eq. **5** and Eq. **6**, $F_{2D}$ is independent of $J_p$, so for fixed $\phi_0$, $F_{2D}$ is minimized with respect to lipid and tether compositions of the membrane ($m$ and $\rho$), which is evaluated numerically (see Fig. S11).

### Locating minima of $F^*(\phi_0)$ from derivatives of $F^*_{3D}(\phi_0)$ and $F^*_{2D}(\phi_0)$

When $f_{obstacle} = 0$, the positions of $F^*(\phi_0)$ minima labeled by the red stars in maintext Fig. $4B - C$ can also be found as the right-most intersections of the derivatives of $F^*_{3D}(\phi_0)$ ($\nabla F^*_{3D}(\phi_0)$) and the negative derivative of $F^*_{2D}(\phi_0)$ (-$\nabla F^*_{2D}(\phi_0)$), where the derivatives of $F^*(\phi_0)$ vanish (Fig. S11B). When $f_{obstacle}$ increase to 0.3, the -$\nabla F^*_{2D}(\phi_0)$ now intersects with $\nabla F^*_{3D}(\phi_0)$ at lower positions, where the spacing between $\nabla F^*_{3D}(\phi_0)$ curves at different $J_p$ values are intrinsically wider (see Fig. S11D). Furthermore, the slope of -$\nabla F^*_{2D}(\phi_0)$ reduces the number of $\nabla F^*_{3D}(\phi_0)$ curves it intersects with, as well as enlarges the horizontal distances between the fewer intersections, contributing to the enhanced selectivity and sensitivity of surface condensation.

### An Intuitive Example Explains the General Principle Behind the Obstacle Effects

Although extracted from the study of surface condensation, the observed principle that the coupled growth of two partners is more sensitive to external driving forces when they grow together is general and should be applicable in a wide range of contexts. To demonstrate this point, we examine one intuitive situation that can occur in tennis practice.

Consider a player practicing tennis with a partner, if the player makes a successful serve (merely over the net and in-bounds) which is also returned successfully by the partner, then we say they have one successful rally. Let's assume that the proficiency of the player ($P$) ranges from 0 to 1, and his serve success rate is given by $R_{serve} = P^{\frac{1}{4}}$. If the partner is a professional coach who returns all successful serve (regardless of serve quality), then the rate of successful rally in the player's practice will be $R_{rally} = R_{serve} \times 1 = P^{\frac{1}{4}}$. On the other hand, if the partner is the player's amateur friend, who can only return high quality serves when $P \geqslant 0.5$, at a serve quality dependent return success rate of $R_{return} = R^{2.9}_{serve}$, then we'll have $R_{rally} = R_{serve} \times R_{return} = P^{\frac{3.9}{4}}$. Here, $R_{serve}$ or $P$ plays the role of external driving force or $J_p$ in surface condensation, $R_{rally}$ the role of $\phi_0$ and $R_{return}$ the role of membrane composition. Indeed we see higher selectivity and sensitivity in the response function of $R_{rally}$ to $P$ when the two partners' performances improve together (see Fig. S18). Although this example is purely imaginary, we hope it helps with the conceptual illustration of the general principle.

### Analysis of the tether-free systems

In this section, we provide more discussions about the GCMC and MFT analysis of the tether-free surface condensation systems presented in the "*General Significance of Membrane-surface polymer Co-operativity to Surface Condensation*" section of the main text.

**A. Varying $h_t$ and $l$ changes the ability of surface polymer to reorganize the underneath membrane.** As discussed in the main text, the membrane's contribution to the formation of surface condensate of density $\phi_0$ as a function of membrane composition is given by main text Eq. **15**. However this expression could also be viewed as the free energy of a membrane at different compositions ($x_A$) when a given amount of surface polymer ($\phi_0$) is present. Thus, when $h_t$ or $D$ ($D$ in MFT essentially represents $l$ in the simulation, see Fig. S5D and S17E) increases, the enthalpic gain of the $f_{int}$ term increases accordingly, and a given $\phi_0$ will shift the minimum of $F_{2D}$ to higher $x_A$ values despite the entropic loss of localizing lipid A (given by the $f_{mem}$ term). Additionally, increasing $D$ also decreases the loss of conformational entropy of confining polymers to the surface phase, which favors the increase of $\phi_0$ and hence $x_A$ (see the third section of the *SI text* above and main text Eq. **4**). Therefore, varying $h_t$ and $D$ changes the ability of surface polymer to reorganize the underneath membrane without perturbing the intrinsic tendency of the membrane to phase separate.

From another perspective, as stated in the section "*Physical Origin of the Obstacle Effects*" in the main text, the formation of surface condensation consists of four contributions 1. the confinement of polymers from a dilute bulk solution into the surface condensate; 2. formation of favorable polymer-polymer interactions in the surface condensate; 3. concentration of specific lipids to the membrane beneath the condensate; 4. formation of favorable tether-polymer interactions in the surface condensate (see main text for details). Surface condensate emerges only when the free energy gain from contributions 2 and 4 outweighs the loss of contributions 1 and 3. Increasing $D$ or $h_t$ then increases the enthalpic gain of contribution 4, which means a smaller enthalpic gain from contribution 2 is required to drive the formation of a certain condensate (labeled by $\phi_0$ and $x_A$). In other words, this means that when $D$ or $h_t$ increases, a smaller $J_p$ is now required to generate a condensate of a given $\phi_0$ (main text Eq. **3** and Eq. **15**), or a condensate of smaller $\phi_0$ (which is favorable for the sum of contributions 1 and 2 as we are studying the prewetting scenario) can be obtained at the same $J_p$, diminishing selectivity. This provides a complementary discussion about selectivity of surface condensation to that provided in the first paragraph of the main text section "*Physical Origin of the Obstacle Effects*".

**B. Free energy analysis of the tether-free system with MFT.** The MFT for the tether-free system differs from the MFT introduced for studying "obstacle effect" only by the removal of tether contributions (see the second to last section of *Methods* in the main text). Hence the method of free energy analysis for tether-free system stays the same as introduced before. Varying $h_t$, $J_m$, $D$ and $\lambda_m$ in the MFT models the effect of varying $h_t$, $J_m$, $l$ and $f_A$ in the GCMC simulation (main text Fig.5$A-H$). Here $h_t$ and $J_m$ have the same meaning in simulation and MFT, $D$ closely represent $l$ as discussed in part A, and $\lambda_m$ and $f_A$ both represent the abundance of lipid A in the membrane. As shown in Fig. S14, in regime I of the parameter space (Fig. S14A-D), $\phi^\infty$ glocally minimizes $F^*(\phi_0)$ and the system shows no prewetting transition. In regime III (Fig. S14E-H), the $\phi_0$ that minimizes $F^*(\phi_0)$ is small yet larger than $\phi^\infty$ at the low $J_p$ value, which gradually becomes larger as $J_p$ increases, reflecting the low selectivity and sensitivity of condensate regulation in this regime. In regime II (Fig. S14I-K), minimization of $F^*(\phi_0)$ at a $\phi_0$ larger than $\phi^\infty$ only happens at large $J_p$ and $\phi_0$ values, reflecting the high sensitivity and selectivity of condensate regulation in this regime.

**C. Additional discussion of the 2D phase diagram in main text Fig.5$J$.** As discussed in the main text Fig.5$J$, $h_t$ and $J_m$ together determine the boundary of the 2D phase diagram that separates the three regimes of surface condensation, corroborating the general principle about condensate regulation presented in our work. Additionally, a noticeable feature of this 2D phase diagram is that, the boundary that separates regime I and II is always sharper than that separates regime II and III, and the sensitivity always reach its highest (darkest color) at the former and gradually diminishes moving towards the latter. Such an observation could be understood by resorting to the free energy analysis of the surface condensation system (Fig. S14). In regime III (Fig. S14E-H), the system is at a condition favorable for surface condensation, (small entropic loss of contribution 3 and large enthalpic gain of contribution 4). In this case, all $J_p$ values enable the formation of a local minimum in $F^*(\phi_0)$ lower than the dilute reference at $\phi^\infty$, indicating low selectivity for surface condensation. Moreover, all these minima are between $\phi^\infty$ and the rightmost (deepest) minimum in the order of $J_p$ they correspond to, indicating a low sensitivity of condensate density to $J_p$. Yet as the parameters of the system are tuned to be less favorable for surface condensation, all local minima of $F^*(\phi_0)$ are elevated gradually, with fewer and fewer remaining below the dilute reference, leading to higher selectivity. Additionally, this also results in the enlarged spacing between the fewer remaining minima between $\phi^\infty$ and the deepest minimum, indicating higher sensitivity and that the system is in regime II. Finally, as the parameters are tuned even more unfavorable for surface condensation, we arrive at the boundary of regime II and I (Fig. S14I-K), where only the rightmost minimum is left below the dilute reference, and we achieve the highest sensitivity and selectivity.

The above observation could also be understood from the perspective of co-operativity. In regime III, a small $\phi_0$ at the lowest $J_p$ could effectively reorganize the membrane and induce surface condensation, which means the condensate density then grows alone slowly as $J_p$ further increases. Moving into regime II, the $\phi_0$ and $J_p$ required to reorganize the membrane become larger (higher selectivity), and co-operative growth of polymer density and local membrane domain is achieved before such $\phi_0$ is reached, enlarging the range of co-operative growth and hence sensitivity. Accordingly, as the system further moves to the boundary of regime II and I, the largest selectivity and range of co-operative growth is observed.

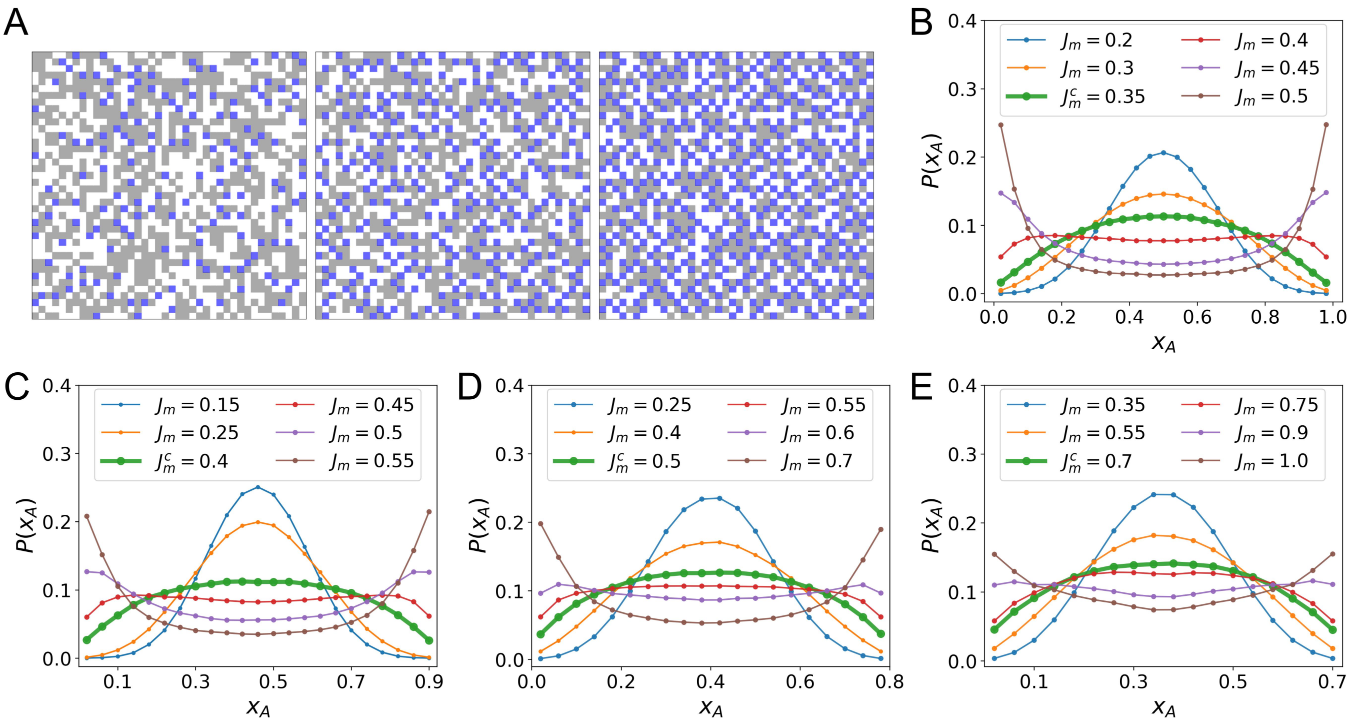


**Fig. S1.** Protein obstacles suppress the critical temperature of membrane. (A) Snapshots of Ising membranes where grey and white squares represent two lipid components A and B, and blue squares the protein obstacles at area fractions of 0.1 (Left), 0.2 (Middle) and 0.3 (Right). (B) Lipid component probability distribution functions $P(x_A)$ for surface regions of $5 \times 5$ lattice sites with different $J_m(k_BT)$ values for membrane without obstacles and membranes with obstacles at area fractions of 0.1 (C), 0.2 (D) and 0.3 (E). $X_A$ is calculated as the number of lipid A sites in regions of $5 \times 5$ divided by 25. The centers of the $P(x_A)$ curves in (C-E) gradually shift to smaller $x_A$ values compared with those of (B), which center at $x_A = 0.5$, is because of the increase in obstacle occupancy.

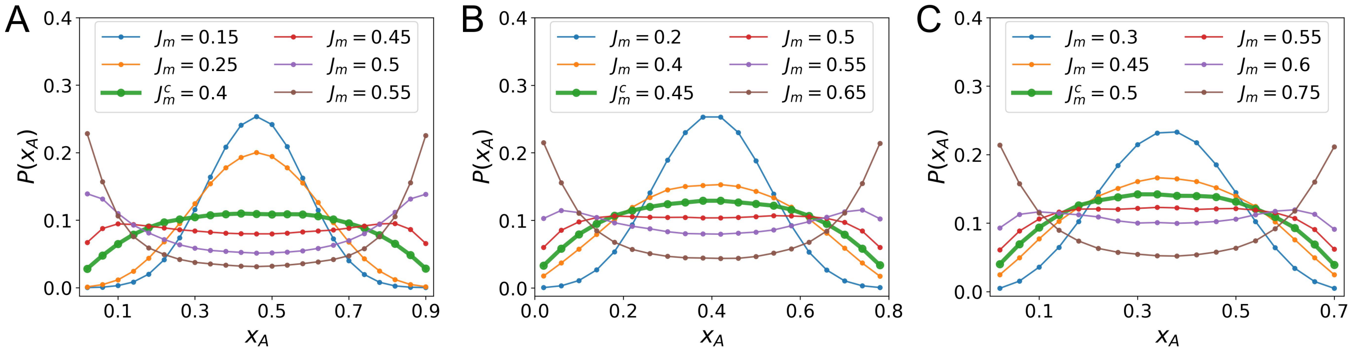


**Fig. S2.** Effect of floating obstacles on membrane critical point. Lipid component probability distribution function $P(x_A)$ for surface regions of 5×5 lattice sites at different $J_m(k_BT)$ for membranes with floating obstacles at area fractions of 0.1 (A), 0.2 (B) and 0.3 (C).

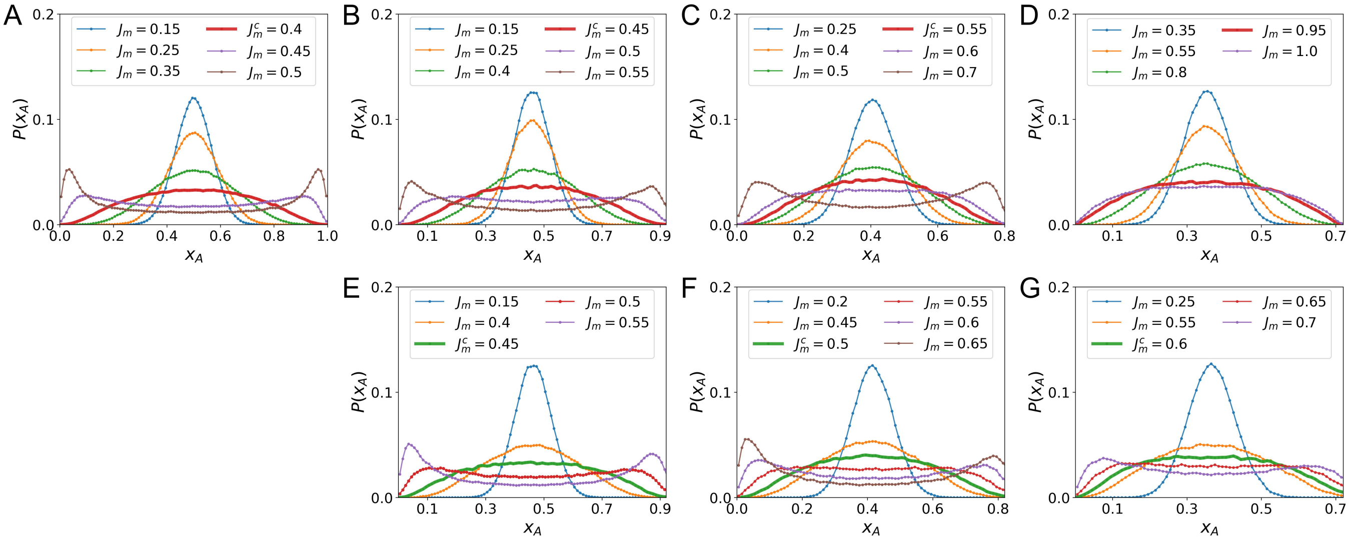


**Fig. S3.** Effect of obstacles on the critical point of larger surface region. Lipid component probability distribution functions $P(x_A)$ for surface regions of 10×10 lattice sites at different $Jm(k_BT)$ for membrane without obstacles (A), membranes with fixed obstacles at area fractions of 0.1 (B), 0.2 (C) and 0.3 (D), and membranes with floating obstacles at area fractions of 0.1 (E), 0.2 (F) and 0.3 (G).

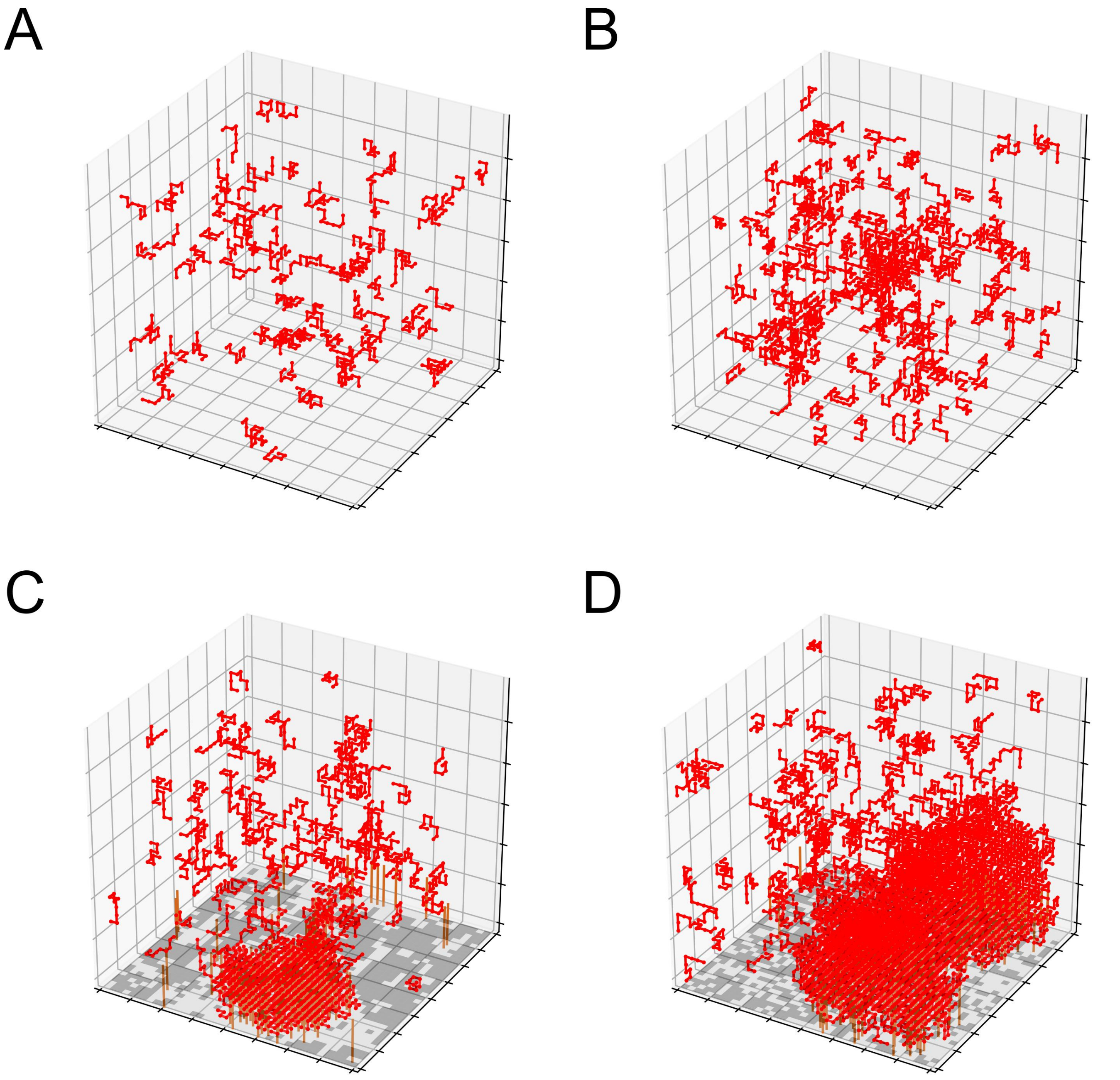


**Fig. S4.** Bulk phase separation, wetting and prewetting. Snapshots of bulk simulation at $J_p$ = 0.5 $k_BT$ (A) and $J_p$ = 0.55 $k_BT$ (B). Bulk phase separation starts to occur at $J_p$ = 0.55 $k_BT$. Snapshots of simulation at $J_m$ = 0.35 $k_BT$ with $J_p$ = 0.5 $k_BT$ (C) and $J_p$ = 0.55 $k_BT$ (D). At $J_p$ = 0.55 $k_BT$, polymer condensates wet the membrane surface, forming surface condensates with finite thickness. When $J_p$ is reduced to 0.5 $k_BT$, where no stable bulk polymer condensate exists, the surface condensate formed becomes a prewetting phase which is molecularly thin.

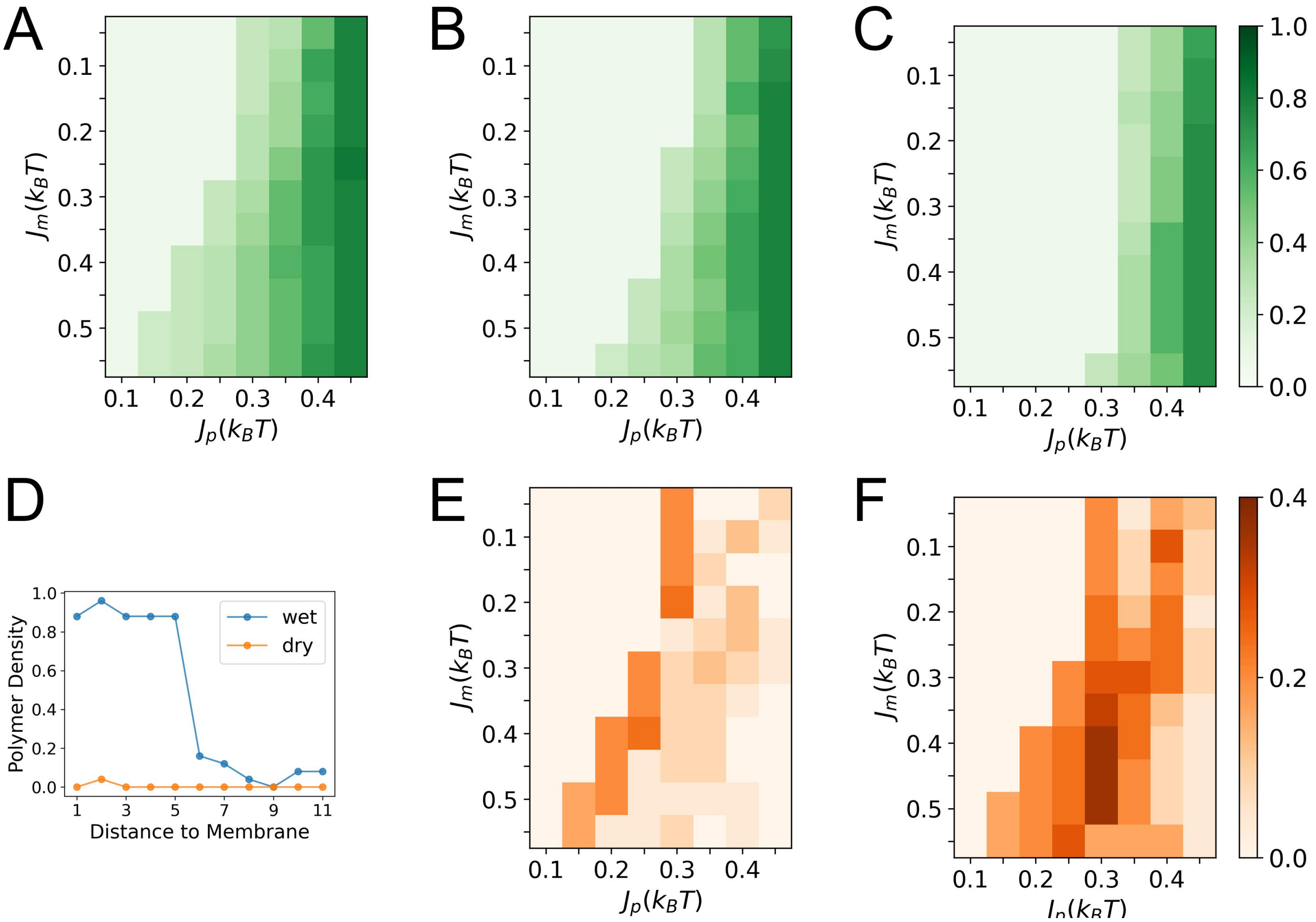


**Fig. S5.** Effect of obstacles on densities of surface condensates. (A) Polymer density in surface condensates formed at different $J_p(k_BT)$ and $J_m(k_BT)$ without obstacles and with obstacles at area fractions of 0.1 (B) and 0.2 (C). (D) Polymer density as a function of distance to membrane at different surface regions of maintext Figure 2C. Blue and orange curves show polymer density profiles of surface regions with and without condensate formation. (E-F) Polymer density differences between (B-C) and (A). Polymer density is calculated as the fraction of lattice sites occupied by red polymers in a $5 \times 5 \times 5$ surface region. The low polymer densities in (A)-(C), (e.g., the columns of $J_p$ = 0.1 $k_BT$) represent the density of dilute surface phase before prewetting transitions.

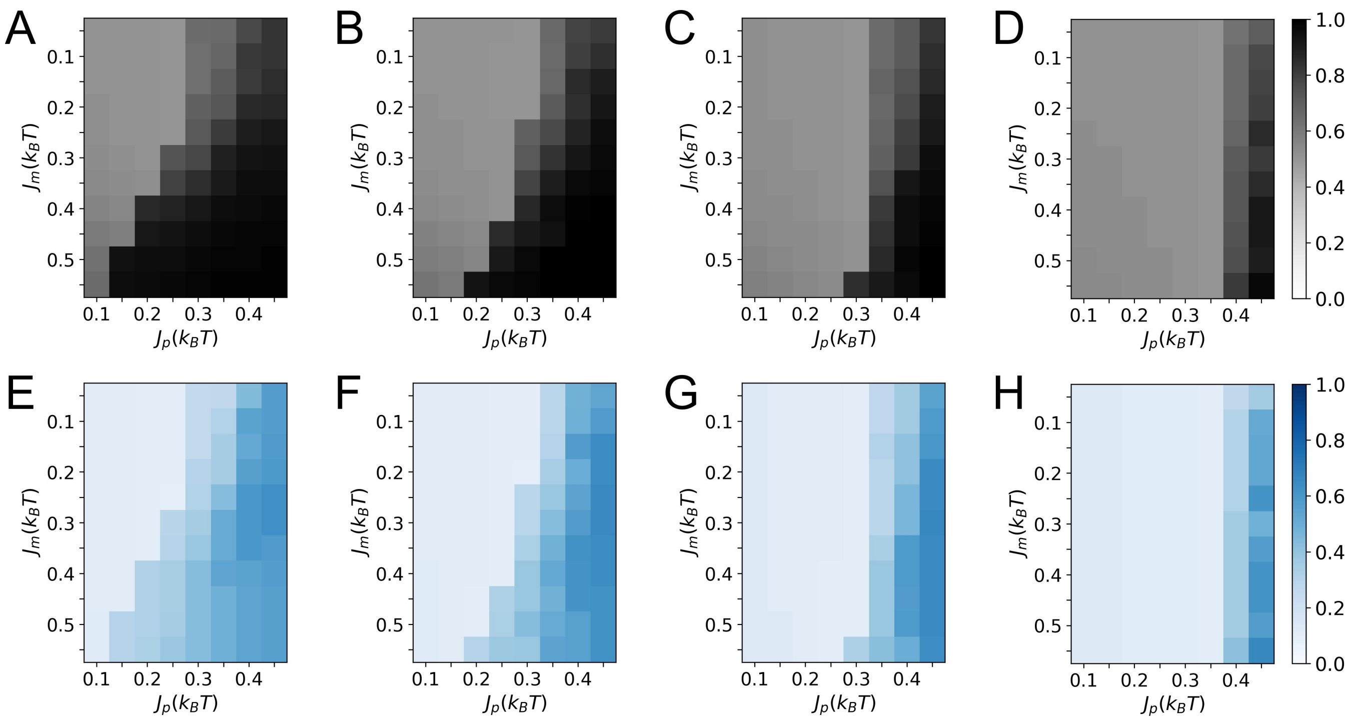


**Fig. S6.** Effect of obstacles on the membrane compositions beneath the surface condensates. (A) Fraction of type-A lipid in all lipids beneath the surface condensates formed at different $J_p(k_BT)$ and $J_m(k_BT)$ without obstacles and with obstacles at area fractions of 0.1 (B), 0.2 (C) and 0.3 (D). (E) Fraction of tethers beneath the surface condensates formed at different $J_p(k_BT)$ and $J_m(k_BT)$ without obstacles and with obstacles at area fractions of 0.1 (F), 0.2 (G) and 0.3 (H). Regions corresponding to low polymer densities in Fig. S5A-C and maintext Fig. 2B, (e.g., the columns of $J_p$ = 0.1 $k_BT$) represent the membrane compositions beneath the dilute surface polymer phase before prewetting transitions.

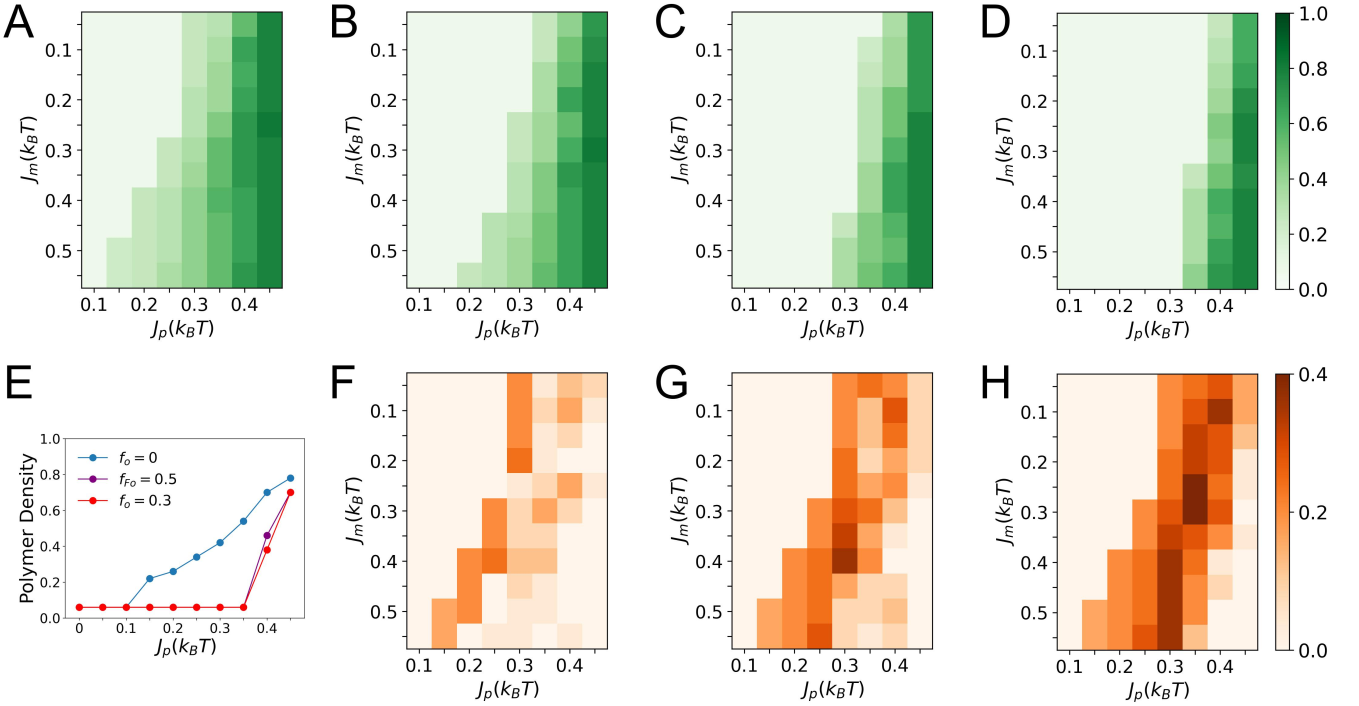


**Fig. S7.** Effect of floating obstacles on the densities of surface condensates. (A) Polymer density in surface condensates formed at different $J_p(k_BT)$ and $J_m(k_BT)$ without obstacles and with floating obstacles at area fractions of 0.1 (B), 0.2 (C) and 0.3 (D). (F-H) Polymer density differences between (B-D) and (A). Polymer density is calculated as the fraction of lattice sites occupied by red polymers in a $5 \times 5 \times 5$ surface region. (E) Polymer density of surface condensate as a function of $J_p$ at fixed $J_m$ = 0.55 $k_BT$ and $\rho_t = 0.2$ with different obstacle components. $f_{Fo}$ $(f_o)$ denotes the fraction of floating (fixed) obstacles. Data for polymer density below 0.1 describe the stable dilute surface polymer phase before the prewetting transition.

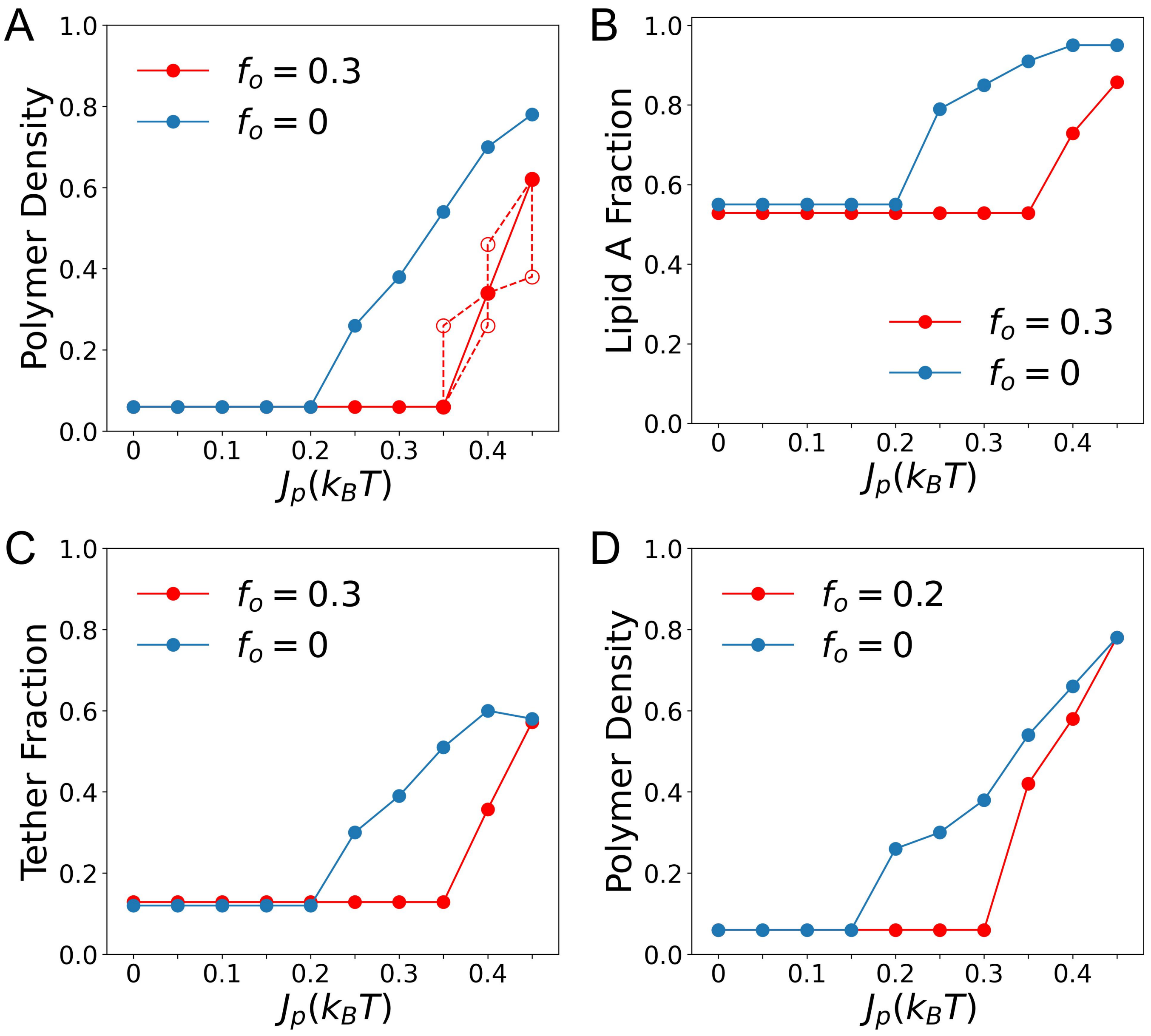


**Fig. S8.** Membrane obstacles enhance the sensitivity and selectivity of surface condensation and its complementary membrane reorganization to $J_p$. (A) Polymer density of surface condensate as a function of $J_p$ at fixed $J_m$ = 0.35 $k_BT$ and $\rho_t = 0.2$. The empty red circles show data for $f_o = 0.3$. Specifically, the empty red circles at $J_p = 0.4\ k_BT$ (lower one) and $J_p = 0.45\ k_BT$ show condensate densities achieved when $J_p$ is increased from 0.35 to 0.4 $k_BT$ and from 0.4 to 0.45 $k_BT$ respectively with membrane configurations frozen. For example, in the simulation corresponds to the empty red circle at $J_p = 0.45\ k_BT$, while $J_p$ is set to 0.45 $k_BT$, the membrane is fixed at representative membrane configuration selected from the trajectory at $J_p = 0.4\ k_BT$. Similarly, the empty red circles at $J_p = 0.4\ k_BT$ (upper one) and $J_p = 0.35$ $k_BT$ show condensate densities obtained when $J_p$ is set to 0.4 and 0.35 $k_BT$ while the membrane is fixed at representative membrane configuration selected from the trajectory at $J_p$ = 0.45 and 0.4 $k_BT$, respectively (B) Fraction of type-A lipid (up spin) in all lipids beneath the surface condensate as a function of $J_p$ at fixed $J_m$ = 0.35 $k_BT$ and $\rho_t = 0.2$. (C) Fraction of tethers beneath the surface condensate as a function of $J_p$ at fixed $J_m$ = 0.35 $k_BT$ and $\rho_t = 0.2$. $f_o$ represent the area fraction of membrane obstacles. (D) Polymer density of surface condensate as a function of $J_p$ at fixed $J_m$ = 0.3 $k_BT$ and $\rho_t = 0.2$ before and after introducing 20 percent obstacles. Unlike other simulations for studying the obstacle effect, the membrane here has three lipid components instead of two. Specifically, the lipid composition is 45 percent lipid A, 45 percent lipid B, and a third inert lipid species at a fraction of 0.1. Data for polymer densities below 0.1 describe the stable dilute surface polymer phase before the prewetting transition.

 

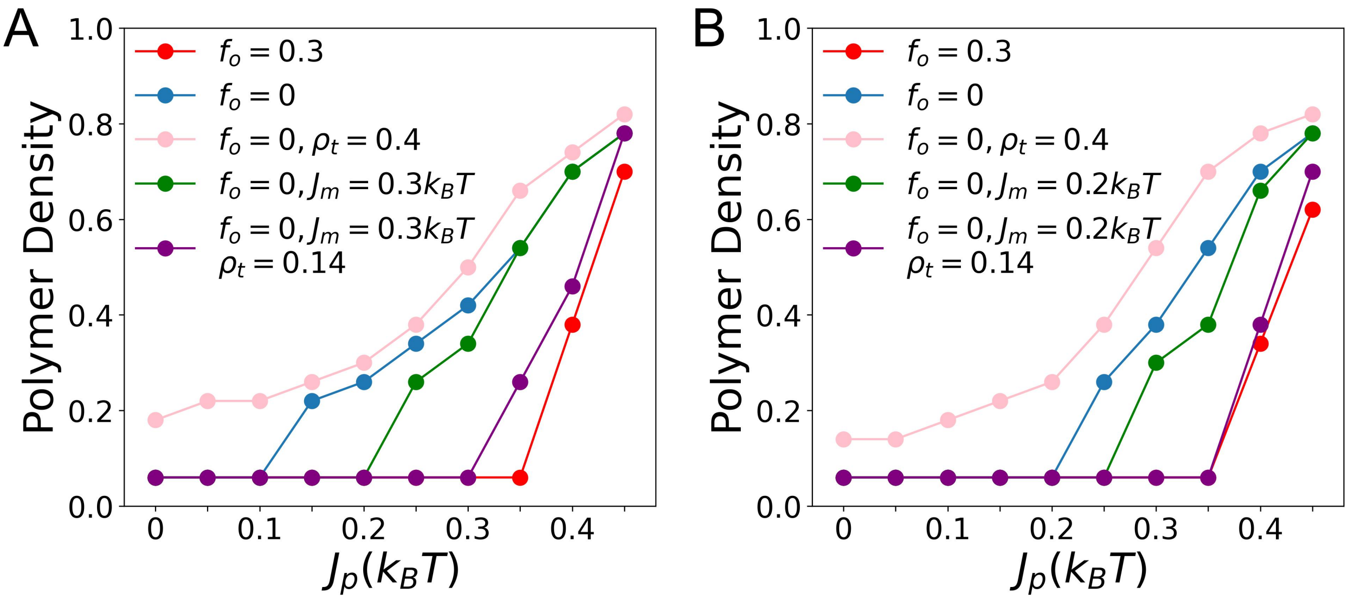


**Fig. S9.** (A) Polymer density of surface condensate as a function of $J_p$ at fixed $J_m$ = 0.55 $k_BT$ and $\rho_t = 0.2$ unless otherwise stated. (B) Polymer density of surface condensate as a function of $J_p$ at fixed $J_m$ = 0.35 $k_BT$ and $\rho_t = 0.2$ unless otherwise stated. Data of low polymer density (<0.1) describe the stable dilute surface polymer phase before the prewetting transition.

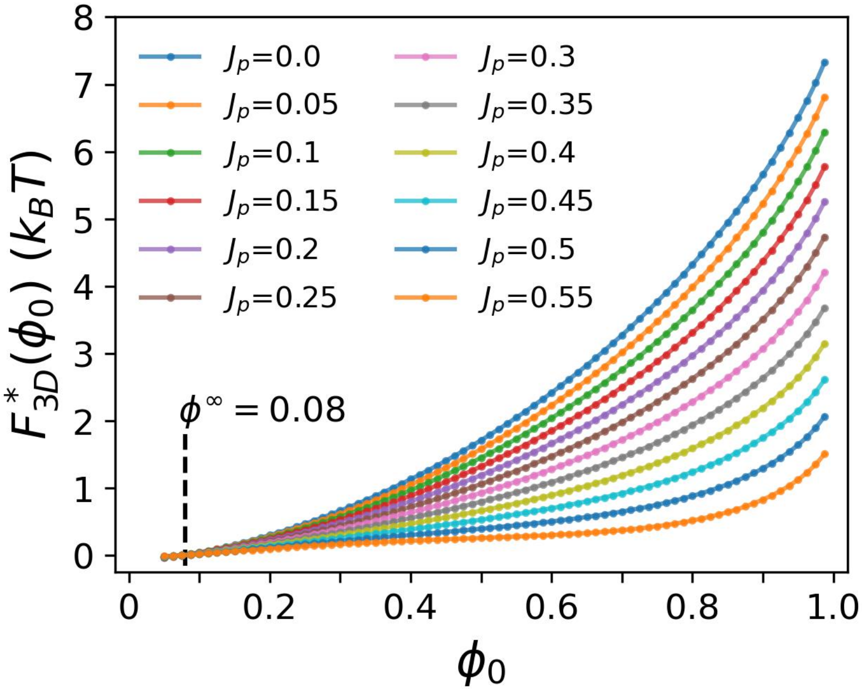


**Fig. S10.** Minimized $F_{3D}$ as a function of polymer density in the surface condensate $F^*_{3D}(\phi_0)$ at different $J_p(k_BT)$.

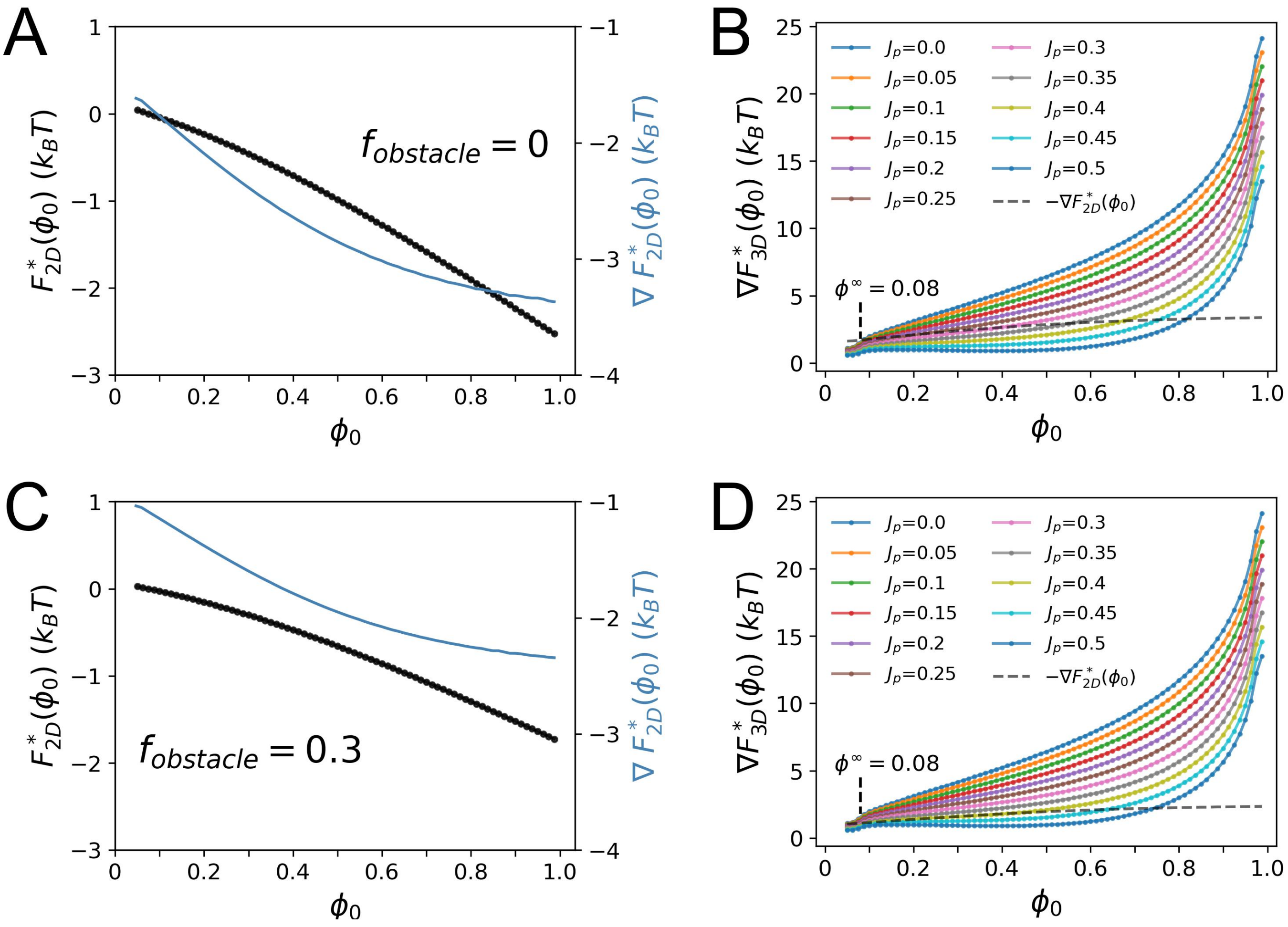


**Fig. S11.** (A) Minimized $F_{2D}$ as a function of polymer density in the surface condensate $F^*_{2D}(\phi_0)$ (black curve, left y axis) and its gradient with respect to $\phi_0$ ($\nabla F^*_{2D}(\phi_0)$) (blue curve, right y axis) at $f_{obstacle}$ = 0. (B) Gradient of $F^*_{3D}(\phi_0)$ with respect to $\phi_0$ ($\nabla F^*_{3D}(\phi_0)$) at different $J_p(k_BT)$ (colored curves) plotted against the negative gradient of $F^*_{2D}(\phi_0)$ (black dashed curve) at $f_{obstacle}$ = 0. (C-D) Same as (A-B) but at $f_{obstacle}$ = 0.3.

**Table S1. Parameters used in the MFT study of obstacle effects.** [a]

| $f_{obstacle}$ | D | $J_m$ ($k_BT$) | $\lambda_m$ ($k_BT$) | $\lambda_\rho$ ($k_BT$) | $h_t$ ($k_BT$) | K ($k_BT$) |
|---|---|---|---|---|---|---|
| 0 | 5 | 0.2 | -0.05 | -0.05 | 0.7 | 1 |
| 0.3 | 5 | 0.2 | -0.05 | -0.05 | 0.7 | 1 |

a. $J_m$ of 0.2 $k_BT$ is close to the critical value of 0.25 $k_BT$ in a obstacle free 2D Ising membrane in the mean-field theory. This is further manifested in Fig. S21.

 

**Table S2. Simulation parameters used in maintext Fig. 5**

| Figure | $J_m$ ($k_BT$) | $f_A$ | $h_t$ ($k_BT$) | $l$ |
|---|---|---|---|---|
| Fig.5$A$ | / | 0.1 | 0.5 | 5 |
| Fig.5$B$ | 0.35 | / | 1.0 | 5 |
| Fig.5$C$ | 0.25 | 0.1 | / | 5 |
| Fig.5$D$ | 0.3 | 0.1 | 1.0 | / |

**Table S3. MFT parameters used in maintext Fig. 5**

| Figure | $J_m$ ($k_BT$) | $\lambda_m$ ($k_BT$) | $h_t$ ($k_BT$) | $D$ |
|---|---|---|---|---|
| Fig.5$E$ | / | -0.24 | 0.36 | 5 |
| Fig.5$F$ | 0.35 | / | 0.5 | 5 |
| Fig.5$G$ | 0.3 | -0.24 | / | 5 |
| Fig.5$H$ | 0.3 | -0.2 | 0.7 | / |

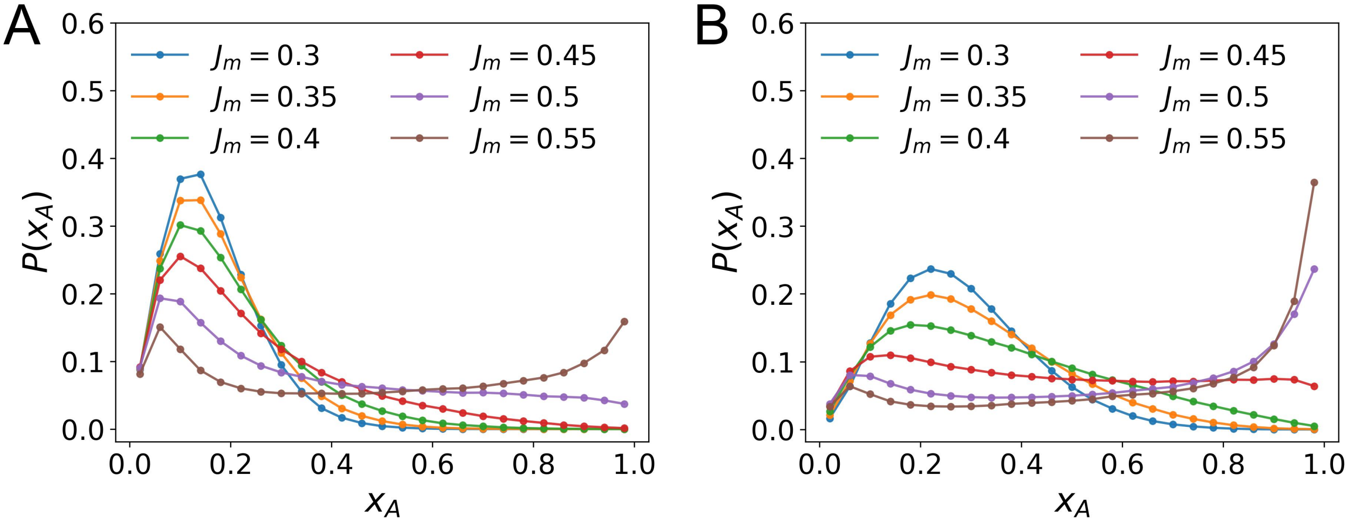


**Fig. S12.** Lipid component probability distribution functions $P(x_A)$ for surface regions of $5 \times 5$ lattice sites with different $J_m(k_BT)$ values for membrane with lipid A fraction ($f_A$) of 0.1 (A) and 0.2 (B). $x_A$ is calculated as the number of lipid A sites in regions of $5 \times 5$ divided by 25. When $f_A$ =0.1 (0.2), $P(x_A)$ is unimodally distributed at low $J_m$ values with a peak centered around $x_A =$ 0.1 (0.2), indicating a well mixed membrane. As $J_m$ increases to the critical value of 0.45 (0.35) $k_BT$, the peak gradually broadens, and when $J_m$ further increases beyond the critical value, $P(x_A)$ becomes bimodally distributed, which indicates the coexistence of two membrane phases with different $x_A$. Increasing $f_A$ of the membrane obviously favors membrane phase separation (when $f_A < 0.5$).

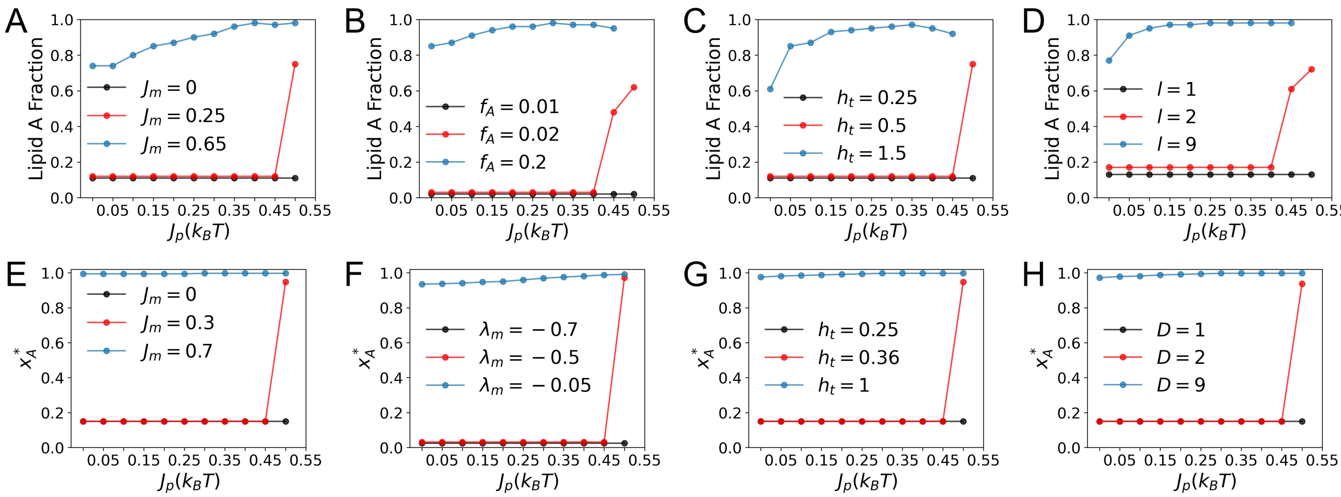


**Fig. S13.** (A-D) Lipid compositions beneath the surface phases shown in maintext Fig.5$A - D$ from GCMC simulations. (E-H) Lipid compositions beneath the surface phases shown in maintext Fig.5$E - H$ from MFT calculations.

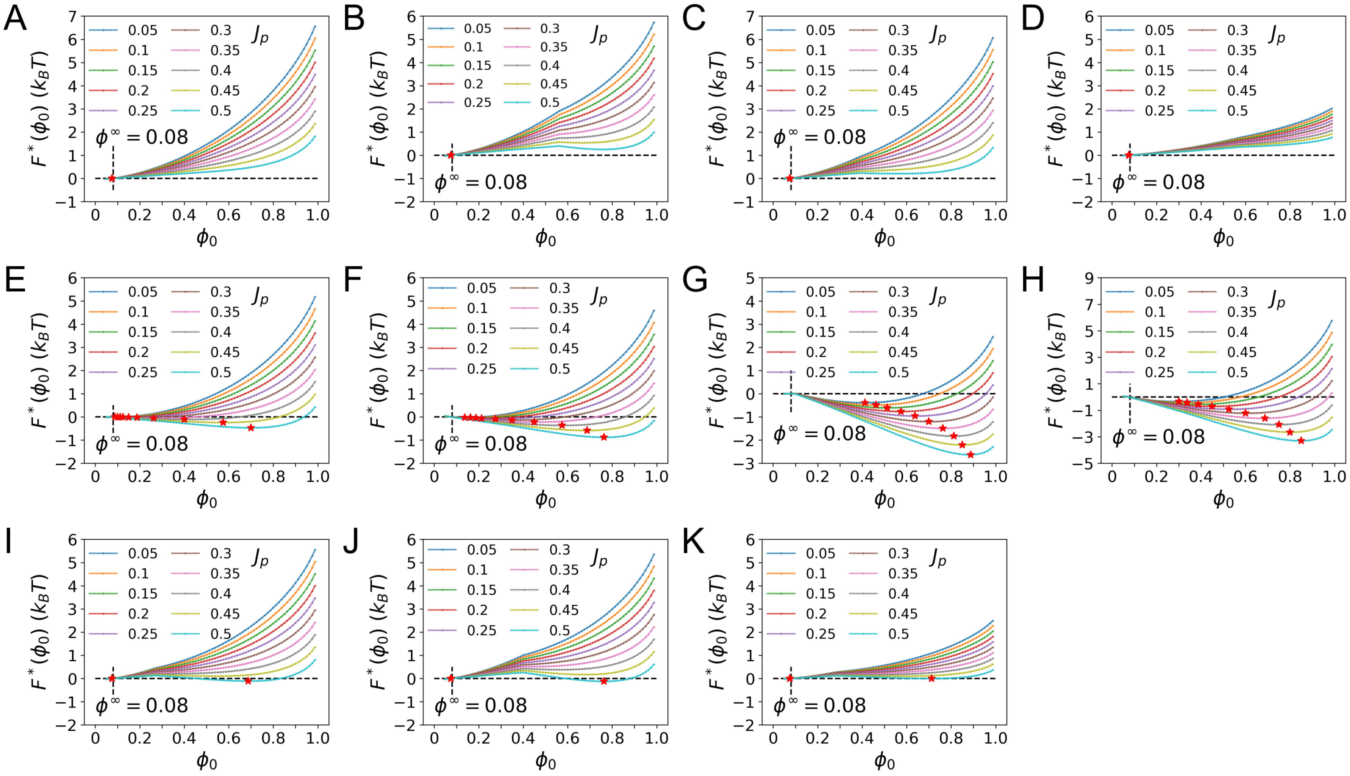


**Fig. S14.** Free energy analysis of the tether-free system from MFT calculations. Minimized free energy as a function of surface polymer density $F^*(\phi_0)$ at different $J_p(k_BT)$ with $\lambda_m = -0.24\ k_BT$, $h_t = 0.36\ k_BT$, $D = 5$ and $J_m =$ (A) 0; (E) 0.7 $k_BT$; (I) 0.3 $k_BT$; corresponding to the data of maintext Fig. 5$E$. $F^*(\phi_0)$ at different $J_p(k_BT)$ with $J_m = 0.35\ k_BT$, $h_t = 0.5\ k_BT$, $D = 5$ and $\lambda_m =$ (B) -0.7 $k_BT$; (F) -0.05 $k_BT$; (J) -0.5 $k_BT$; corresponding to the data of maintext Fig. 5$F$. $F^*(\phi_0)$ at different $J_p(k_BT)$ with $J_m = 0.3\ k_BT$, $\lambda_m = -0.24$, $D = 5$ and $h_t =$ (C) 0.25 $k_BT$; (G) 1 $k_BT$; (I) 0.36 $k_BT$; corresponding to the data of maintext Fig. 5$G$. $F^*(\phi_0)$ at different $J_p(k_BT)$ with $J_m = 0.3\ k_BT$, $\lambda_m = -0.2$, $h_t = 0.7\ k_BT$ and $D =$ (D) 1; (H) 9; (K) 2; corresponding to the data of maintext Fig. 5$H$. Red stars label the positions where $\phi_0$ minimizes $F^*(\phi_0)$ at different $J_p(k_BT)$.

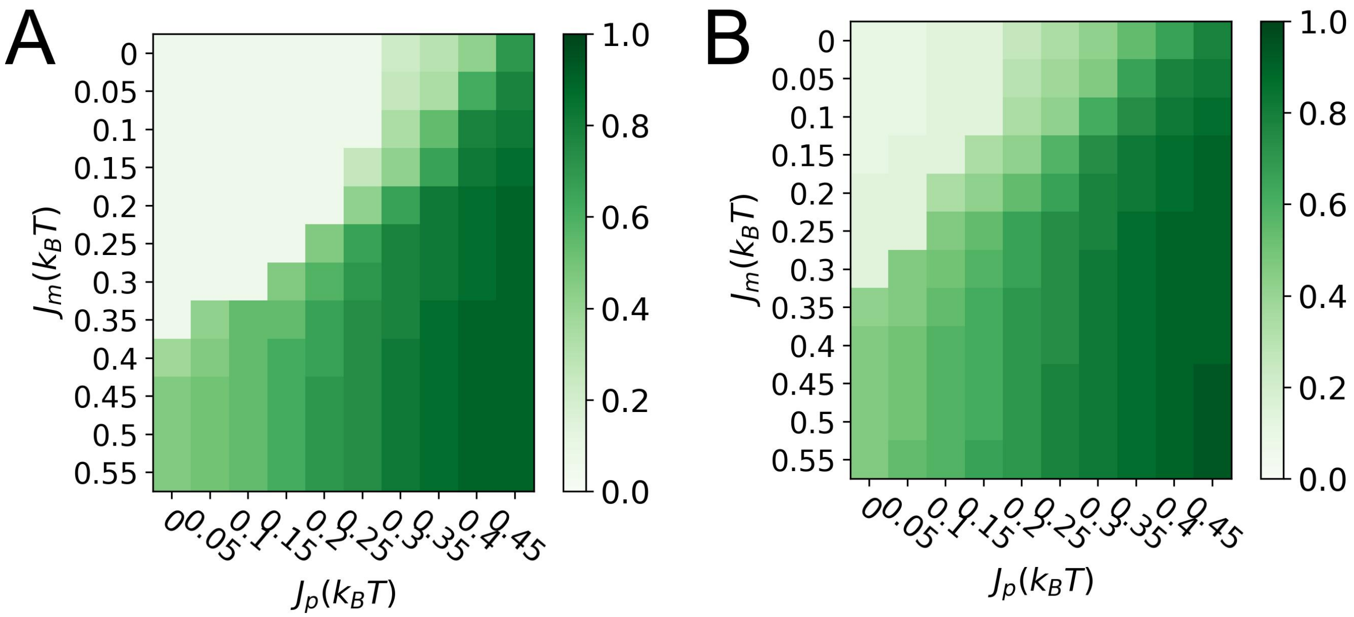


**Fig. S15.** Effect of $f_A$ on surface condensate density. GCMC result of surface condensate density at different $J_p(k_BT)$ and $J_m(k_BT)$ values with $h_t = 1\ k_BT$, $l = 5$ and (A) $f_A = 0.1$; (B) $f_A = 0.2$. Data of low polymer density (<=0.1) describe the stable dilute surface polymer phase before the prewetting transition.

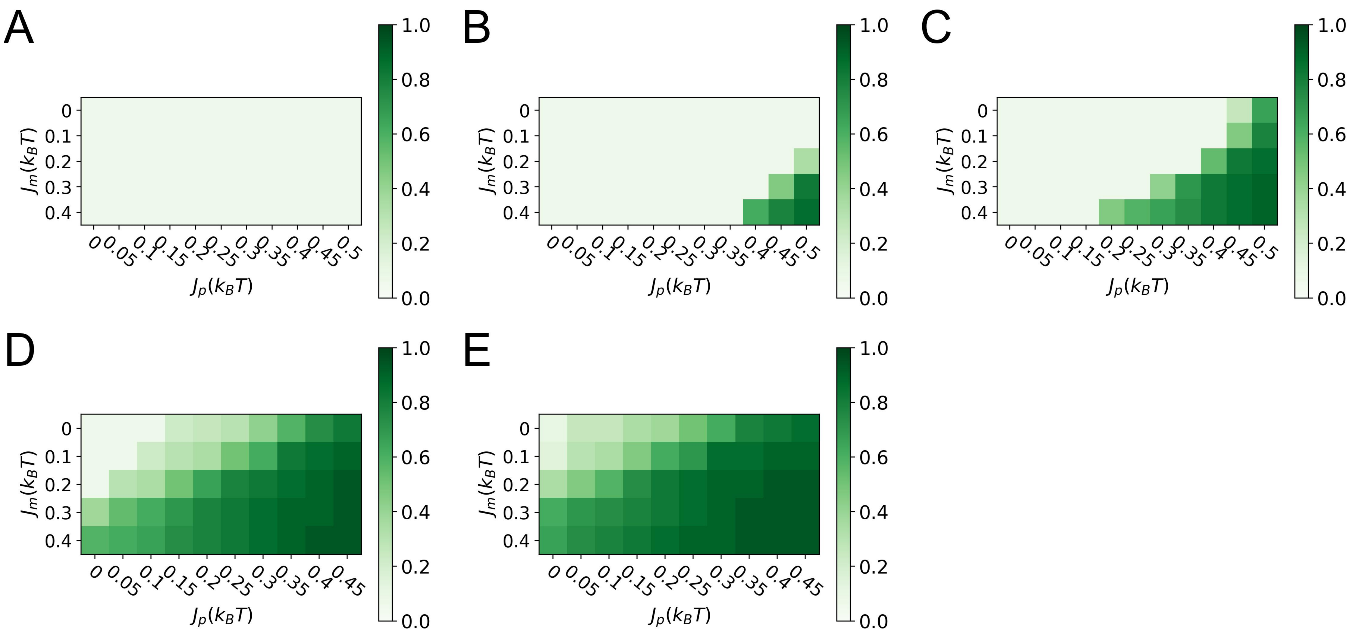


**Fig. S16.** Effect of $h_t$ on surface condensate density. GCMC result of surface condensate density at different $J_p(k_BT)$ and $J_m(k_BT)$ values with $f_A = 0.1$, $l = 5$ and (A) $h_t = 0.25\ k_BT$; (B) $h_t = 0.5\ k_BT$; (C) $h_t = 0.75\ k_BT$; (D) $h_t = 1.25\ k_BT$; and (E) $h_t = 1.5\ k_BT$. Data of low polymer density (<0.1) describe the stable dilute surface polymer phase before the prewetting transition.

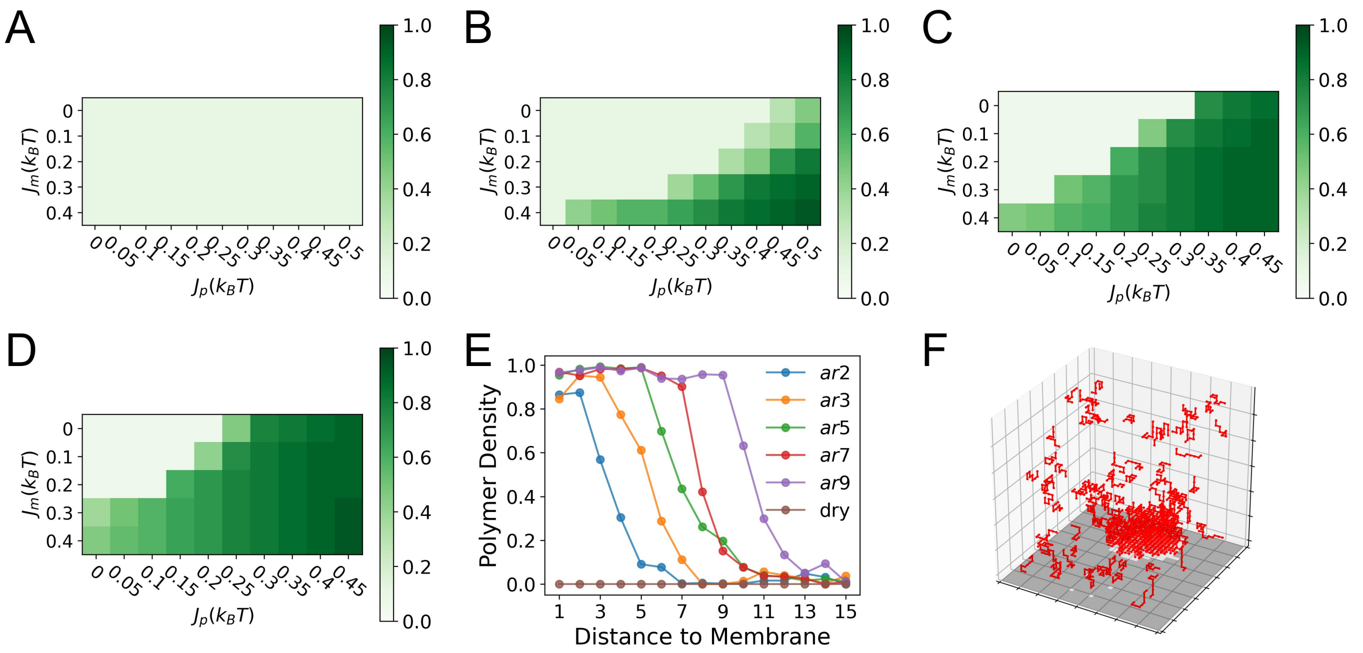


**Fig. S17.** Effect of $l$ on surface condensate density. GCMC result of surface condensate density at different $J_p(k_BT)$ and $J_m(k_BT)$ values with $f_A = 0.1$, $h_t = 1\ k_BT$ and (A) $l = 1$; (B) $l = 3$; (C) $l = 7$; and (D) $l = 9$; Data of low polymer density (<0.1) describe the stable dilute surface polymer phase before the prewetting transition. (E) Effect of $l$ on the thickness of surface condensate. (F) a snapshot of the GCMC simulation with $J_p = 0.45\ k_BT$, $J_m = 0.4\ k_BT$, $f_A = 0.1$, $h_t = 1\ k_BT$ and $l = 5$.

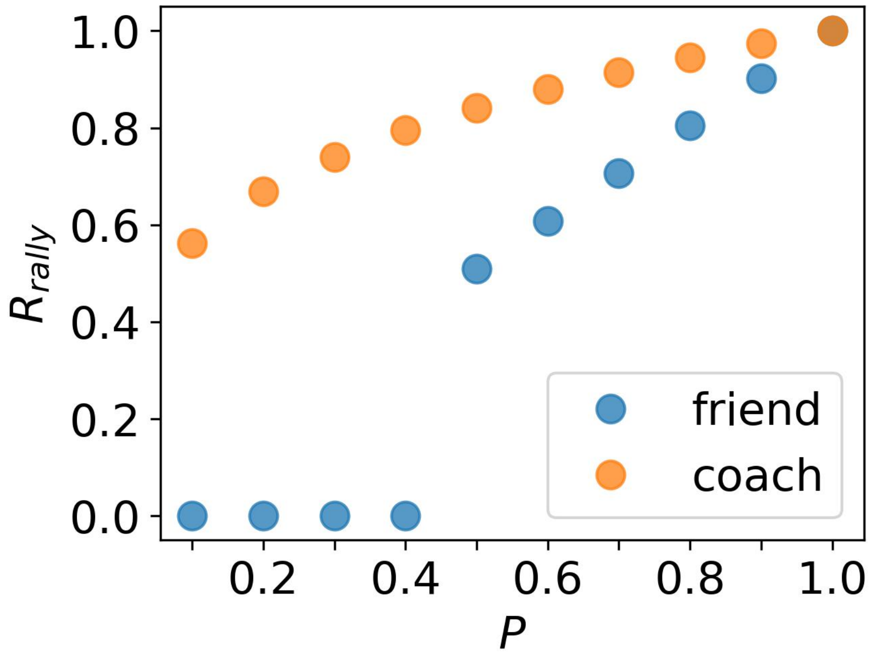


**Fig. S18.** Intuitive example from tennis explaining the general principle that enhancement of sensitivity to external driving force originates from coupled growth of two order parameters.

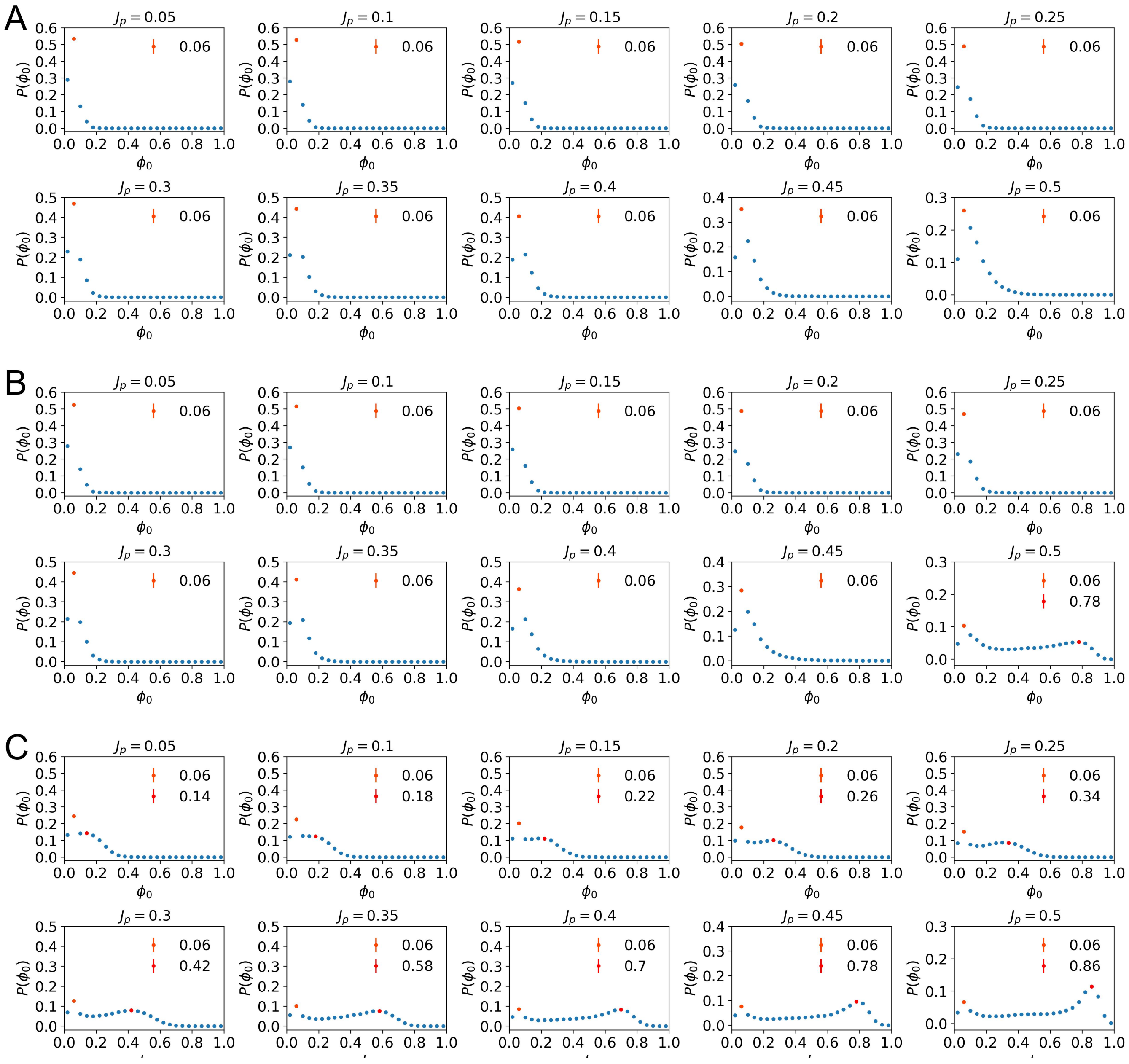


**Fig. S19.** Identification of surface condensation and condensate density in GCMC study. Surface polymer density distribution at $J_p$ from 0.05 to 0.5 $k_BT$ with $f_A = 0.1$, $h_t = 0.5\ k_BT$, $l = 5$ and $J_m =$ (A) 0; (B) 0.25 $k_BT$ and (C) 0.65 $k_BT$. (A-C) corresponds to the black, red and blue data of maintext Fig.5$A$. Peak locations are found by finding local maxima of the raw data or data after background subtraction (e.g., subtract the data of $J_p = 0.45\ k_BT$ from that of $J_p = 0.5\ k_BT$ in (B) for finding the location of the second peak in the latter). Error bars show the s.e.m. of the data which are usually smaller than the data label size.

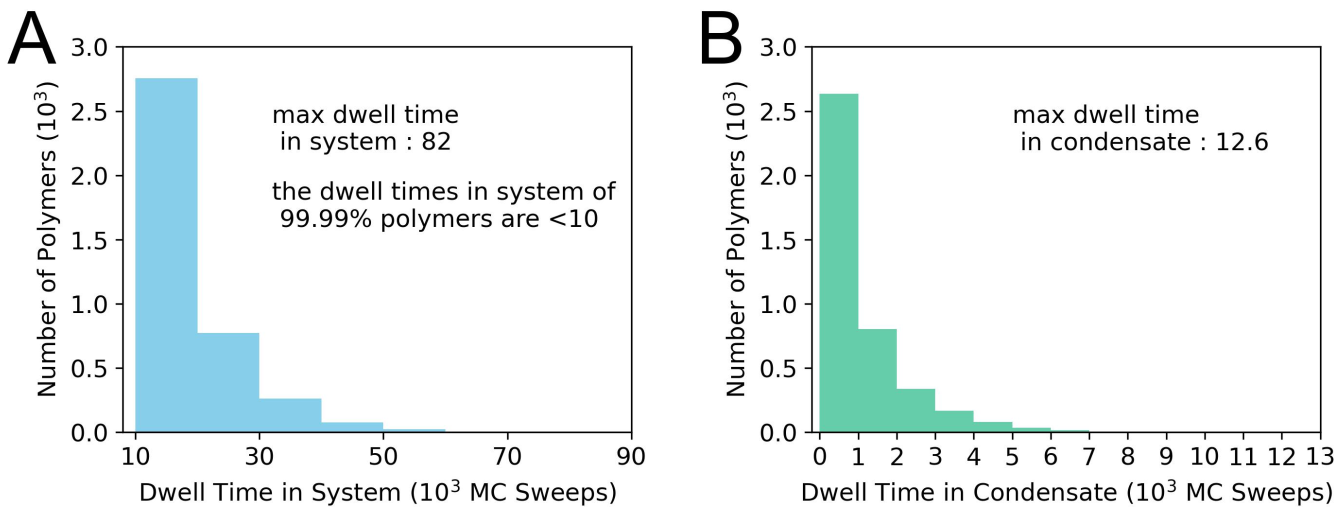


**Fig. S20.** Dwell time distribution of polymers in the simulation system (A) and surface condensate (B). The data are from the simulation at $J_p = 0.45\ k_BT$, $J_m = 0.55\ k_BT$ and $f_{obstacle} = 0$, where the density of surface condensate reaches the highest value among all simulation conditions and consequently results in worst polymer exchange efficiency between condensate and the dilute solution. Still, the typical dwell time of a polymer chain in the condensate is below 6000 MC sweeps, which is more than 2 orders of magnitude smaller than the simulation time (1 million MC sweeps).

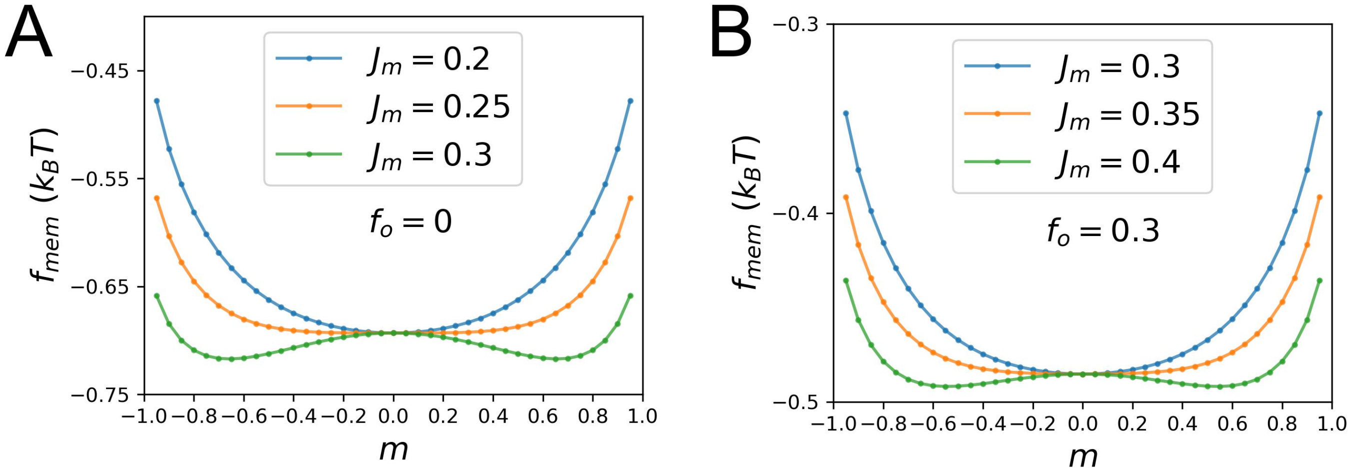


**Fig. S21.** Membrane free energy (Eqn. **6** of the main text with $\lambda_m = 0$) at different $J_m$ values with obstacle fractions of 0 (A) and 0.3 (B).
